# An ITK – SLP-76 Y173 Axis Regulates TCR Signal Persistence and Thymic Development

**DOI:** 10.64898/2026.10.22.760889

**Authors:** Manal Marzouk, Jillian K. Perry, Enas Hallumi, Hirofumi Shibata, Zenia Kaul, Naama Klopstock, Dominic P. Golec, Leslie J. Berg, Pamela L. Schwartzberg, Deborah Yablonski

## Abstract

The TCR-induced tyrosine kinase cascade results in ITK-mediated phosphorylation of at least two substrates: PLC-γ1 Y783, an essential mediator of TCR signaling, and SLP-76 Y173. We previously provided evidence that SLP-76 Y173 supports proximal TCR and FcεRI signaling; but its role in thymocyte development was never explored. Here we demonstrate that SLP-76 Y173 was required for TCR/CD28-induced degradation of IκB; moreover, both ITK and SLP-76 Y173 were required for persistent TCR-induced phosphorylation of PLC-γ1, ERK and p38. To explore whether SLP-76 Y173 may mediate ITK-dependent events in vivo, we replaced SLP-76 Y173 with phenylalanine in a mouse model. Like ITK-deficient thymocytes, SLP-76 Y173F thymocytes exhibited reduced thymic cellularity, reduced CD4/CD8 ratios, and reduced surface expression of CD5. In ex vivo-stimulated thymocytes, mutation of SLP-76 Y173 or inhibition of ITK reduced the persistence of TCR-induced calcium flux; moreover, TCR-induced phosphorylation of ERK and p38 depended on SLP-76 Y173. Like ITK-deficient thymi, SLP-76 Y173F thymocytes exhibited decreased production of CD4^−^ iNKT_1_ thymocytes, coupled with increased production of innate-like CD4^+^ and CD8^+^ SP thymocytes. Our results provide strong evidence that the ITK substrate site, SLP-76 Y173, promotes TCR signal persistence that is required to mediate the thymic developmental effects of ITK.

**One Sentence Summary:** An ITK substrate site, SLP-76 Y173, is a predominant mediator of the signaling and developmental effects of ITK.

**Key Points:**

- An ITK substrate site, SLP-76 Y173, regulates TCR signal amplitude and persistence
- Persistent TCR-induced phosphorylation of p38 depends on ITK and SLP-76 Y173
- The thymic developmental effects of ITK are predominantly mediated by SLP-76 Y173

## Introduction

Thymocyte development is tightly regulated by signaling through the T cell antigen receptor (TCR). TCR ligation indirectly triggers the assembly of a large signaling complex, often referred to as the LAT signalosome, that is nucleated by three adaptors: SLP-76, Gads and LAT [1]. Signalosome assembly is triggered by a TCR-activated, Syk-family tyrosine kinase, ZAP-70, which phosphorylates four tyrosines on the cytoplasmic tail of LAT and three at the N-terminus of SLP-76 [2, 3]. Further downstream, a Tec-family tyrosine kinase, ITK, is activated within the signalosome, where it exerts profound effects on TCR signaling and thymocyte development [4].

A central role of ITK is the activation of PLC-γ1. ITK phosphorylates PLC-γ1 at two sites that are required for its activation, Y783 and Y775 [5–8]. Active PLC-γ1 produces inositol triphosphate (IP3) and diacylglycerol (DAG), second messengers that initiate downstream transcriptional and developmental responses. Consistent with this paradigm, ITK-deficient peripheral T cells show reduced PLC-γ1 phosphorylation and impaired calcium signaling [5].

A complex network of protein-protein interactions is required to bring about ITK-mediated phosphorylation of PLC-γ1. PLC-γ1 is recruited to the signalosome by its N-terminal SH2 domain, which binds to phospho-LAT Y132 (Y136 in mouse) [9, 10]. ITK may bind bivalently to SLP-76, as its SH2 domain binds to SLP-76 p-Y145, while its SH3 domain binds an adjacent proline-rich motif [11–13], which may also bind the SH3 domain of PLC-γ1 [14, 15]. Binding of ITK to SLP-76 maintains the active conformation of ITK, while facilitating ITK-mediated phosphorylation of LAT-bound PLC-γ1 [8].

Additional ITK substrates within the signalosome include SLP-76 Y173 and Gads Y45 [16, 17]. Outside of the signalosome, ITK may phosphorylate the transcription factors T-bet and TFII-I [18, 19]. Just as ZAP-70 directly phosphorylates multiple essential targets within the TCR signaling pathway, we considered the possibility that multiple ITK substrates may be required to mediate its profound signaling and developmental functions.

We first identified p-Y173 in the course of a phospho-mass spectrometry analysis of AspN-cleaved SLP-76, as this site was not detectable following trypsin cleavage [16]. TCR-and FcεRI-induced phosphorylation of Y173 were validated using a phospho-specific antibody [16].

Functional analyses, conducted in a Jurkat-derived human T cell line and in bone-marrow-derived mast cells, demonstrated that SLP-76 Y173 is required to support TCR-induced PLC-γ1 phosphorylation, calcium flux and NF-κB activation, as well as FcεRI-induced phosphorylation of p38 [16, 17].

Further supporting the importance of an ITK-SLP-76 Y173 signaling axis, auxiliary expression of membrane-tagged SLP-76 substantially increases the antigen sensitivity of CAR T cells; remarkably, this activity absolutely depended on the ITK-recruitment site, SLP-76 Y145, and the ITK substrate site, SLP-76 Y173 [20]. Yet the mechanisms by which SLP-76 Y173 interfaces with downstream responses are not yet understood. Here, we provide evidence that an ITK-SLP-76 Y173 axis is essential to mediate the biological effects of ITK in the TCR signaling pathway and in thymic development.

Conventional thymocyte development proceeds through discrete stages that are strictly regulated by TCR signaling. Within the CD4^−^CD8^−^(DN) compartment, basal pre-TCR signaling triggers β-selection, characterized by upregulation of CD5 and passage of DN3 cells through DN4 and onto the CD4^+^CD8^+^ (DP) compartment [21, 22]. At this stage, TCR-mediated recognition of self-peptide-MHC (p-MHC) antigens on thymic epithelial cells triggers cell fate decisions that are associated with further upregulation of CD5 and transient upregulation of CD69 [22, 23]. Whereas high affinity p-MHC recognition triggers negative selection and clonal cell death, moderate-affinity recognition triggers positive selection, accompanied by differentiation and passage of thymocytes to the CD4^+^ or CD8^+^ SP compartments [24, 25].

As central components of the signalosome that mediates pre-TCR and TCR signaling, SLP-76 and LAT are absolutely required for thymocyte development beyond the DN stage [26–28]. ITK^−/−^ mice exhibit milder thymic phenotypes. Thymic cellularity is moderately decreased, and the ratio of CD4 to CD8 SP cells is reduced [29, 30]. A subtle, pre-TCR signaling defect impairs the upregulation of CD5 at the DN to DP transition [30, 31]. A defect in positive selection was revealed when ITK^−/−^mice were crossed onto certain TCR transgenic backgrounds [29, 30, 32]. Together, these phenotypes are consistent with partial impairment of pre-TCR and TCR signaling.

In addition to its role in conventional thymocyte development, ITK is required to support the development of invariant natural killer T cells (iNKT) [33–35], an innate-like, agonist-selected lineage that recognizes CD1d-presented lipid antigens [24, 36]. Consistent with the mouse phenotype, ITK-deficient patients lack NKT cells and exhibit profoundly increased susceptibility to EBV [37].

While promoting iNKT cell development, ITK suppresses the development of innate-like SP populations that express characteristic transcription factors: eomesodermin (EOMES) or promyelocytic leukemia zinc finger (PLZF). PLZF is a key regulator of innate-like T-cell differentiation that promotes the acquisition of innate effector functions [38, 39]. EOMES is highly expressed in memory-like CD8 SP thymocytes and is required for their development and characteristic rapid effector responses [40, 41]. These populations functionally interact, as IL-4 produced by PLZF^+^ thymocytes induces EOMES expression and promotes the development of innate, memory-like CD8 SP thymocytes [40, 42].

ITK^−/−^ thymi include a large, memory-like, EOMES^+^ CD44^hi^ CD122^+^ CD8 SP population, known for its ability to rapidly produce IFN-γ [42, 43]. The CD4 SP compartment also features an unusual EOMES^+^ CD4 SP population [41], and expanded populations of PLZF^+^ CD4 thymocytes that may express either an αβ or γδ TCR [42, 44]. Overall, these findings show that ITK is required to regulate the balance between conventional and innate-like thymocyte development.

Multiple lines of evidence suggest that the distinctive phenotypes of ITK^−/−^ thymocytes derive from a loss of ITK kinase activity. Many of the ITK^−/−^ phenotypes are recapitulated in SLP-76 Y145F mice, which express ITK, but lack the SLP-76 site responsible for recruiting and activating ITK [40, 41, 45]. Moreover, selective ablation of the ITK kinase domain suffices to induce expansion of a memory-like CD8 SP population [46]. Together, these findings support the notion that ITK kinase activity profoundly influences thymocyte development, yet the ITK substrates responsible for limiting the expansion of innate-like subsets have not been identified. Here, we examine the downstream signaling functions of a recently identified ITK substrate, SLP-76 Y173, and provide evidence that this site is required to mediate signaling and developmental functions of ITK.

## Materials and Methods

### Antibodies

Anti-mouse fluorescently conjugated antibodies CD4-PerCP-Cy5.5, or -FITC, or –PE (Clone RM4-4 and GK1.5), CD8a-APC-Cy7, or –FITC, or –PE (Clone 53-6.7 or QA17A07), TCR β chain-BV605, or PE-Cy7 (Clone H57-597), CD44-BV421 (Clone IM7), CD5-BV711 (Clone 53-7.3), CD69-BV605 (Clone H1.2F3), CD122 (IL-2Rβ)-FITC, or –PE (Clone TM-β1), CD62L-BV605 (Clone MEL-14), TCR *γ*/δ-PerCP-Cy5.5 (Clone GL3), CD25-APC (Clone PC61), CD3ε -APC (Clone 145-2C11), TCR Vβ8.1,8.2-PE-Cy7 (Clone MR5-2), TCR Vβ5.1,5.2-PE-Cy7 (Clone MR9-4), CD45R/B220-FITC (Clone RA3-6B2), TER-119/Erythroid Cells-FITC (Clone TER-119), CD11b-FITC (Clone M1/70), EOMES-AF647 (Clone W17001A), PLZF-PerCP-Cy5.5 (Clone 9E12), T-bet-BV605 (Clone 4B10), Zombie Aqua Fixable Viability Kit (423101), Isotype control antibodies Rat IgG2a,κ-FITC (Clone RTK2758), Rat IgG2b,κ-FITC (Clone RTK4530), anti-IκB-PE (clone 3D6C02), anti-p38 MAPK Phospho (Thr180/Thr182)-PE (Clone A16016A), anti-PLC*γ*1 Phospho (Tyr783)-PE (Clone A17025A), anti-LAT Phospho (Thy171)-AF647 (Clone A20005D), anti-ERK1/2 Phospho (Thr202/Tyr204)-AF647 (Clone 4B11B69), biotin-conjugated anti-mouse antibodies CD3ε (Clone 145-2C11), CD4 (Clone RM4-4), and CD8 (Clone 53-6.7), were from BioLegend. Indo-1 AM Calcium Sensor Dye from eBioscience™. Anti-mouse fluorescently conjugated antibodies CD4-BUV737 (Clone GK1.5), CD8α-BUV395 (Clone 53-6.7), CD24-BUV496 (Clone M1/69), TCR β chain-BV650 (Clone H57-597), CD44-BV786 (Clone IM7), TCR *γ*δ-BV711 (Clone GL3), ROR *γ*t-PE-CF594 (Clone Q31-378), streptavidin-PE, and anti-mouse FITC-conjugated antibodies used for TCR Vβ screening panel were from BD Biosciences. Anti-mouse biotin-conjugated antibody CD122 (Clone TM-b1), and fluorescently conjugated antibodies CD5-APC-eFlour 780 (Clone 53-7.3), and PLZF-AF488 (Clone Mags.21F7) were from eBioscience. Live/Dead Aqua-BV510 (L34966) from Invitogen. PBS-57 loaded and unloaded CD1d-tetramers were from NIH Tetramer Core Facility.

### Cell Lines

All cell lines used in this study derive from J14 [47], a SLP-76-deficient derivative of the Jurkat T cell line. As previously described [17], J14 cells were retrovirally reconstituted with wild type or mutant, N-terminally FLAG- or twin-strep-tagged SLP-76, using an IRES-GFP tagged retroviral vector. Reconstituted cells were sorted for comparable expression of GFP followed by verification of comparable SLP-76 expression, as previously shown [17]. Prior to use, we periodically validate comparable expressions of TCR and CD28 on all cell lines to be used in experiments.

### Mice

Mice were maintained and treated under specific pathogen free (SPF) conditions in accordance with the approved protocols of Animal Care and Use committees of NHGRI (protocol G98-3) and NIAID (protocol LISB-22E) at the NIH (Animal Welfare Assurance #A-4149-01), and in accordance with the Technion Institutional Animal Care and Use Committee ethics approval numbers IL-170-11-22, IL-171-11-22 and IL-062-04-24. All authors were trained and certified to conduct experiments in rodents at the Technion or at the NIH.

All mice were on the C57BL/6 background. ITK-deficient (ITK^−/−^) mice were previously described [29]. To create the SLP-76 Y173F mice we employed CRISPR/Cas9-mediated genome editing to alter the 3’ edge of Lcp2 (SLP-76) exon 7 on mouse chromosome 11, while substituting phenylalanine for tyrosine at codon 173. The gRNA sequence was CTTACCGATGTACATGGAGT and the donor oligo sequence was CTCCAACAACGAGGAGGCTCTGCAGAGCTCCATCCTGCCCCCCAACTCTTTCCACAACAC<u>G</u>AA<u>T</u>TCCATG**<u>TT</u> <u>C</u>**ATCGGTAAGGGCCTCTTGGTAACTTGCAGCCCTGGGAGAAGCTGCTCATTGAGTTTCTCT (The Y to F mutation sequence is bold underlined; simple underline marks silent mutations that inactivate the PAM while creating a new EcoRI site, shown in small caps). Genomic DNA was prepared from potential founder mice, and the region surrounding the mutation was amplified by PCR, tested for sensitivity to EcoRI digestion, and sequenced. Mice derived from two independent founders were bred to homozygosity and found to exhibit similar phenotypes. One of the founders was backcrossed five times onto the C57BL/6 background for in depth analysis. Eight potential off-target sites were sequenced to verify a lack of off-target mutations.

### T cell stimulation and lysis

Jurkat-derived T cell lines were washed and resuspended in stimulation medium (RPMI + 2 μg/ml Glutamine + 20 mM HEPES pH 7.3), preheated to 37°C for 10-15 min, and then stimulated for the indicated times at 37°C with anti-TCR (C305, 1:1000) and anti-CD28 (clone 28.2 2 μg/ml). Cells were lysed in ice-cold lysis buffer (20 mM HEPES pH 7.3, 1% Triton X-100, 0.1% n-dodecyl-β-D-maltoside, 150 mM NaCl, 10 mM NaF, 1 mM Na_3_VO_4_, 1 mM DTT, 10 μg/ml aprotinin, 2 mM EDTA, 10 μg/ml leupeptin, 2 mM phenylmethanesulfonyl fluoride (PMSF), 1 μg/ml pepstatin, and 1 mM dithiothreitol). Thymocytes were collected and lysed by directly adding 2X lysis buffer. Cell lysates were incubated on ice and then centrifuged in a microcentrifuge to pellet cell debris, prior to SDS-PAGE and western blot. Western blots were developed with SuperSignal^TM^ West Pico PLUS Chemiluminescent Substrate, and digital images of the membrane were produced by Fusion Fx7 camera system.

### Long terms stimulation and qPCR

Cells were stimulated for up to 4 hours in RPMI + 1% Penicillin/Streptomycin + 20 mM Glutamine + 10% FCS with plate-bound anti-TCR (C305, 1:10,000) and soluble anti-CD28 (clone 28.2 2 µg/ml), or with PMA (50 ng/ml), Ionomycin (1µM), and anti-CD28 (clone 28.2 2 µg/ml). Cytoplasmic and nuclear extracts were prepared as described [48]. Following stimulation, cells were resuspended in cold buffer A (10 mM HEPES pH 7.9; 10 mM KCI, 0.1 mM EDTA; 0.1 mM EGTA; 1 mM DTT; 0.5 mM PMSF) by gentle pipetting. The cells were incubated on ice for 15 min, after which a 10 % solution of Nonidet NP-40 (Fluka) was added, followed by vortexing. Following cytoplasmic fraction extraction, nuclear extract was resuspended in ice-cold buffer B (20 mM HEPES pH 7.9; 0.4 M NaCl; 1 mM EDTA; 1 mM EGTA; 1 mM DTT; 1 mM PMSF) and incubated on ice for 15 with shaking once every 3 min. The nuclear extract was centrifuged for 15 min in a microfuge at 4 °C. Cytoplasmic and nuclear extracts were analyzed by Western blot. GAPDH and SNF2H were used as cytoplasmic and nuclear loading controls, respectively. mRNA was purified according to the Quick-RNA MiniPrep kit and then converted to cDNA by the iScript cDNA Synthesis Kit. cDNA was diluted as recommended by the standard curve, and the qPCR reaction was performed as indicated in the iTaq Universal SYBR Green kit, using these primers: CAACCGAACATACCCTTCTATCC for c-REL, TCCTGTCTTGCATTGCACTAAG for IL-2, AGTGCAGAAAAACGCCAAGT for Nur77, GGAGCGAGATCCCTCCAAAAT for GAPDH, TGACACTGGCAAAACAATGCA for HPRT1. mRNA was isolated and quantified by qPCR, with normalization to GAPDH and HPRT1.

### Thymocyte isolation and staining for FACS analysis

Thymocytes were suspended in MACS buffer (Ca^2+^/Mg^2+^-free PBS + 2% FBS + 1% Penicillin/Streptomycin + 1 mM EDTA), passed through a 40 µm cell strainer and stained for 30 min at RT with Zombie Aqua Fixable Viability Kit (BioLegend, 423101). In some experiments, thymocytes were suspended in complete medium (RPMI + 10% FBS + 1% Penicillin/Streptomycin + 1% Glutamine + 0.1% BME), passed through a 70 µm strainer and stained at with LIVE/DEAD™ Fixable Aqua Dead Cell Stain Kit (ThermoFisher, L34966). Cells were then washed with FACS buffer (Ca^2+^/Mg^2+^-free PBS + 1-2% FBS and 0.02% sodium azide, with or without 1mM EDTA) and stained for 30 min with fluorescently conjugated antibodies in FACS buffer, at 4°C or at RT if tetramers were included in the staining panel. Where indicated, a mixture of FITC-labeled antibodies was used as a dump gate to exclude non-DN cells; cells that stained positively for CD45R/B220, TER-119/Erythroid Cells, CD11b, as well as CD4 and CD8 expressing cells. Stained cells were washed sequentially with FACS buffer and Ca^2+^/Mg^2+^-free PBS and fixed for 20 min at RT with IC Fixation Buffer (Thermo Fisher Scientific, 00-8222-49) diluted 1:1 in Ca^2+^/Mg^2+^-free PBS, or with 4% PFA (Electron Microscopy Sciences), then washed with FACS buffer. For intracellular TCR detection, fixed cells were washed twice with Permeabilization Buffer (Invitrogen, 00-8333-56), and intracellular staining was performed in the same buffer for 30 min at RT. Prior to transcription factor staining, cells were fixed either for 30-40 min at RT using the True-Nuclear Transcription Factor Buffer Set (BioLegend, 424401), followed by intracellular staining for 30 min RT in Permeabilization buffer, or cells were fixed for 25 min at 4°C using the FoxP3 Transcription Factor Staining Buffer Set (Thermo Fisher Scientific), followed by intracellular staining in FoxP3 Set buffer for a minimum of 1 hour at 4°C.

### Calcium fluorimetry of thymocytes

Thymocytes were barcoded with CellTrace Far Red as described [49], mixed together, and calcium fluorimetry was performed essentially as described [50]. In brief, a mixture of differentially barcoded thymocytes was washed with calcium staining medium (RPMI + 1% FBS + 20mM Hepes pH 7.3 + 2mM glutamine), and incubated for 30 min at 30°C, with 0.5 mL per each 5 million cells of calcium staining medium containing 4 μM indo-1 AM (eBioscience), 2 mM probenecid, fluorescent cell surface markers, including CD4-FITC (clone GK1.5) and CD8-PE (clone QA17A07), and biotinylated stimulatory antibodies: CD3ε (Clone 145-2C11), CD4 (clone RM4-4) and CD8 (clone 53–6.7), 3 μg/mL each. The stained cells were then washed twice and resuspended in stimulation medium (RPMI + 20 mM Hepes pH 7.3 + 2 mM Glutamine). Cells were stored on ice and preheated to 37°C for 10 min prior to ratiometric calcium fluorimetry, which was performed by flow cytometry, with the temperature maintained at 37°C. Stimulation was induced at the 60 s time point by adding streptavidin (Biolegend, 405150) to a final concentration of 25 μg/mL, or ionomycin to a final concentration of 1 μM. Excitation was at 355 nm and emission was measured using the bandpass filters 405/20 (Ca-bound indo-1) and 485/22 (free indo-1), with data presented as the ratio between these two values. Data analysis was performed while gating on DP or SP thymocytes.

### Phospho-flow assays

Murine thymocytes or Jurkat-derived cells were washed with Ca^2+^/Mg^2+^-free PBS, and differentially barcoded using CellTrace Violet, for 20 min at RT with gentle rocking. The staining reaction was terminated by adding 8 mL of stopping medium (RPMI + 20% FBS) and incubating cells for an additional 5 min at RT with gentle rocking. Cells were then combined, washed with stimulation medium (RPMI + 20 mM Hepes pH 7.3 + 2 mM Glutamine), and, in the case of thymocytes, subsequently stained for 30 min at 4°C with CD4-FITC (clone GK1.5), CD8-APC-Cy7 (clone 53–6.7), and 3 μg/mL each of the biotinylated stimulatory antibodies: anti-CD3ε (clone 2C11), anti-CD8 (clone QA17A07), and anti-CD4 (clone RM4-4) in stimulation medium + 1% FBS. Cells were then washed twice, resuspended in stimulation medium, and preheated to 37°C for 15 min prior to stimulation. Thymocytes were stimulated by adding streptavidin to a final concentration of 25 μg/mL. Jurkat-derived cells were stimulated with anti-TCR (C305, 1:1000) with or without anti-CD28 (clone 28.2 2 µg/ml), in the presence or absence of an ITK inhibitor (3µM, BMS-509744). Stimulations were terminated by the addition of an equal volume of IC Fixation Buffer (Thermo Fisher Scientific, 00-8222-49) and incubation for 15 min at 37°C; then cells were washed twice with FACS buffer, followed by permeabilization using pre-chilled True-Phos permeabilization buffer (BioLegend, 425401), added dropwise while vortexing, and incubated for at least 60 min at −20°C, then washed twice with FACS buffer and stained with the phospho-protein antibodies for 30 min at RT.

### Statistical Analysis

All experiments in this study were performed two or more times with similar results. Statistical analysis was performed using the Prism software package. *p*-values were determined using the statistical tests indicated in figure legends. The following symbols were used to indicate statistically significant *p*-values: *p < 0.05; **p < 0.01; ***p < 0.001; ****p < 0.0001

## Results

### TCR/CD28 signaling to NF-κB depends on SLP-76 Y173

We previously showed that Y173 is required for TCR/CD28-induced nuclear translocation of c-Rel, a prominent member of the NF-κB family [17]. To explore whether NF-κB signaling may be more broadly affected, we examined TCR/CD28-induced phosphorylation and degradation of IκBα, an upstream regulator of NF-κB family members.

We evaluated NF-κB in J14-derived cell lines, either lacking SLP-76 or reconstituted with wild type (WT) or Y173F SLP-76. Consistent with a previous report [51], SLP-76 was absolutely required for the TCR/CD28-induced phosphorylation (Fig 1A) and degradation of IκBα (Fig1B-C), which we assessed by using a FACS-based assay to measure the frequency of cells that reduced IκBα. In SLP-76-reconstituted cells, the Y173F mutation almost completely abrogated the ability of SLP-76 to rescue IκBα phosphorylation and degradation (Fig 1A-C), suggesting that this is essential for SLP-76 dependent signaling to NF-κB.

**Figure 1.**
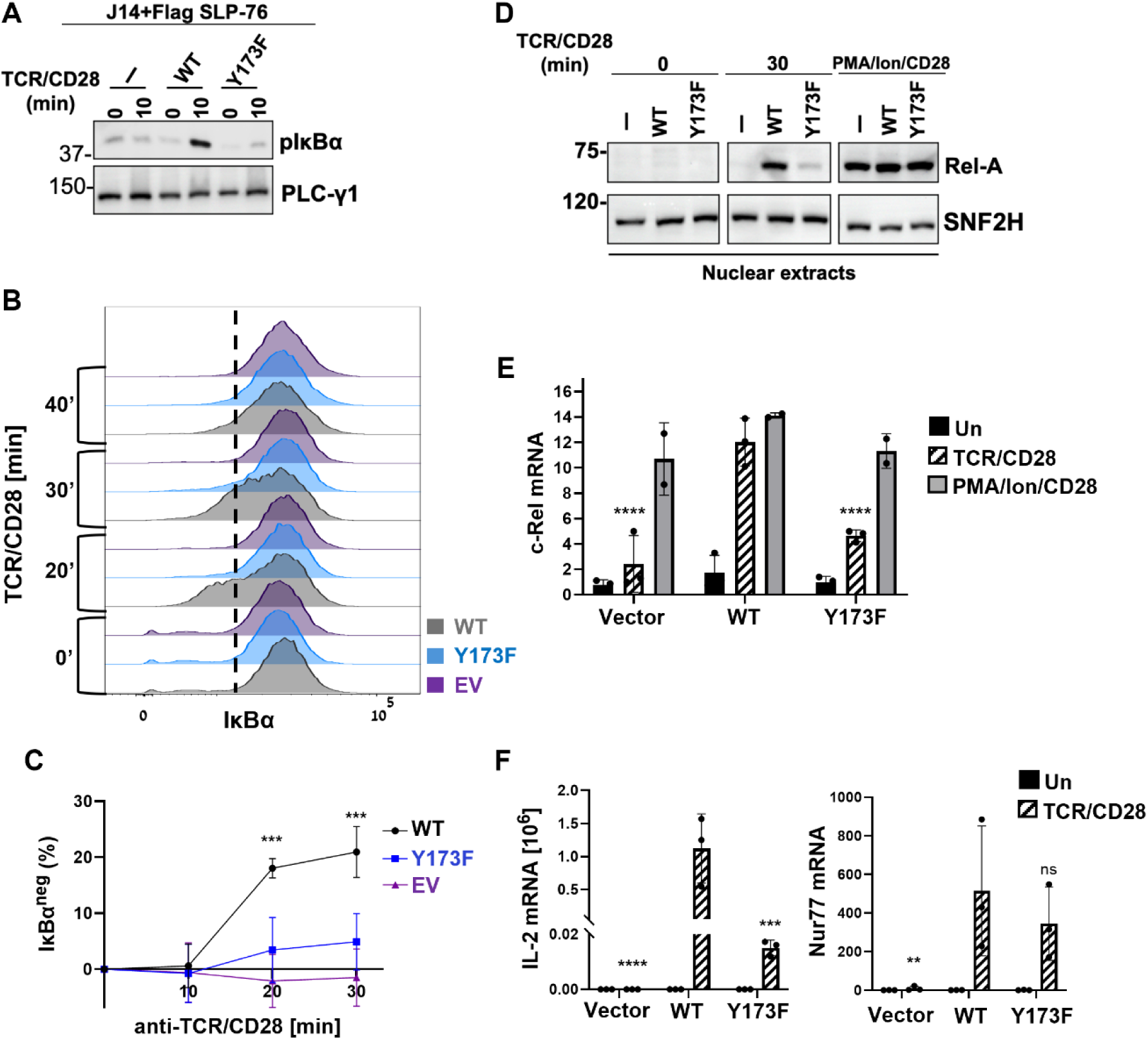
TCR/CD28 signaling to NF-κB depends on SLP-76 Y173. J14 cell lines, stably reconstituted with the indicated alleles of SLP-76, were costimulated with TCR/CD28, prior to assessing cellular responses. (**A**) Following stimulation with soluble anti-TCR/CD28, whole cell lysates were probed by western blotting with the indicated antibodies. Results are representative of two experiments with similar results. (**B-C**) Cells were differentially barcoded with Cell Trace Violet, mixed together and stimulated as in A. At the indicated time points, cells were fixed, permeabilized and stained intracellularly for IκB. Cells were J14, stably reconstituted with WT SLP-76 (grey), Y173F (blue) or empty vector (EV, purple). (**B**) Representative results, the indicated gate was used to define IκB^neg^ cells. (**C**) Average frequency of baseline subtracted IκBα^neg^, data are pooled from three independent experiments; error bars indicate the standard deviation. Statistical significance relates to comparison between Y173F and WT at each time point. (**D**) Cells were stimulated for 30 min with plate-bound anti-TCR, or with PMA and ionomycin, both with soluble anti-CD28, and nuclear extracts were probed by western blotting with the indicated antibodies. Results are representative of three experiments with similar results. (**E**-**F**) Following 4 hours stimulation as in D, qPCR was used to quantify the indicated mRNA, with normalization to GAPDH and HPRT1. Statistical significance relates to comparison between TCR/CD28-stimulated mutant and WT cell lines. Results are the average of three biological repeats with two technical repeats for each. Statistical analyses were by 2-way ANOVA with Dunnett’s correction (**C**), or Tukey’s correction (**E**-**F**).

Further downstream, the Y173F mutation markedly impaired the nuclear translocation of p65/RelA (Fig 1D). TCR/CD28-induced transcription of c-Rel and IL-2 were likewise reduced; yet transcription of Nur77 was not significantly changed (Fig 1E-F). Together, these results confirm that Y173 is essential for TCR/CD28 signaling to NF-κB, while suggesting that at least some aspects of TCR/CD28 signaling may be Y173-independent.

These experiments did not resolve whether Y173 is primarily required for TCR signaling or for CD28 costimulatory activity. To address this question, we next applied phospho-flow to examine the TCR signaling in the absence of CD28 costimulation.

### An ITK–SLP-76 Y173 axis promotes the amplitude and persistence of TCR signaling

#### Y173-dependent phosphorylation of PLC-γ1

We began by assessing the role of SLP-76 Y173 in the TCR-induced phosphorylation of PLC-γ1 Y783, a key regulatory site that is required for PLC-γ1 activation [6]. Both ITK and SLP-76 are required for PLC-γ1 Y783 phosphorylation [5, 17, 47], in part, because SLP-76 is required to recruit and activate ITK, which in turn phosphorylates PLC-γ1 at Y783 [8].

To clarify the role of SLP-76 Y173 in this process, we assessed phospho-PLC-γ1 Y783 on a single cell basis, using phospho-flow. In WT cells, p-PLC-γ1 staining peaked by 2 minutes, followed by a relatively rapid return to just above baseline (Fig 2A). This assay did not produce distinct peaks of responding and non-responding cells, yet we observed reproducible, time resolved changes in the intensity of p-PLCγ1 staining. Further validating the assay, PLC-γ1 phosphorylation was markedly reduced in SLP-76-deficient J14 cells and was nearly abrogated when cells were stimulated in the presence of an ITK inhibitor (Fig 2A).

**Figure 2.**
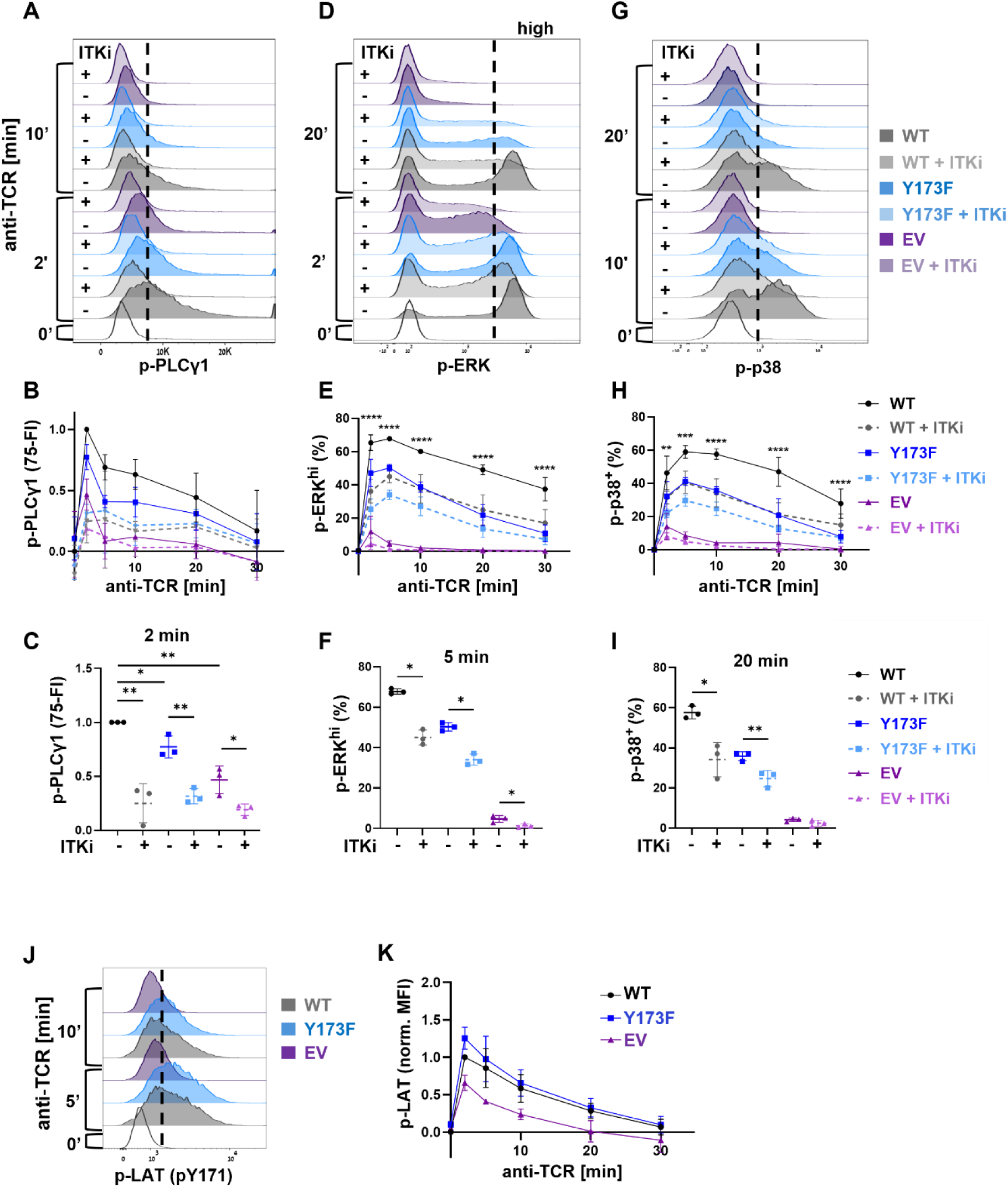
SLP-76 Y173 promotes the amplitude and persistence of TCR signaling. The indicated cell lines were differentially barcoded, mixed together, and incubated for 15 min at 37°C in the presence or absence of an ITK inhibitor (ITKi, BMS-509744, 3 μM), prior to stimulation with soluble anti-TCR (C305, 1:1000). At the indicated time points, cells were fixed, permeabilized, and stained intracellularly for phospho-flow. Cells were J14, reconstituted with WT SLP-76 (grey), Y173F (blue) or empty vector (EV, purple); ITK-inhibited cells are presented in lighter shades. (**A**) Representative data obtained for p-PLCγ1 (Y783). (**B**) Time course: Min/max normalized 75th percentile fluorescence intensity (75-FI) of p-PLCγ1. Zero represents the 75^th^ percentile of WT at time zero, and 1 represents the 75^th^ percentile of WT at 2 min. (**C**) Min/max normalized 75th percentile FI at the 2 min time point, with and without ITKi. (**D**) p-ERK, representative data. (**E**) Time course: Frequency of pERK-^hi^ cells. This gate was set to include 60% of WT cells at the 10 min time point. Statistical significance relates to comparison between WT and Y173F cells, in the absence of ITK inhibition. (**F**) Frequency of pERK^hi^ cells at the 5 min time point, with and without ITKi. (**G**) p-p38, representative data. (**H**) Time course: Baseline-subtracted frequency of p-p38^+^ cells, defined by the gate shown in G. Statistical significance relates to comparison between WT and Y173F cells, in the absence of ITK inhibition. (**I**) Frequency of p-p38^+^ cells at the 20 min time point, with and without ITKi. (**J**) p-LAT (Y171), representative data. (**K**) Time course: Min/max normalized median fluorescence intensitity (MFI) of p-LAT (Y171). Zero represents the WT intensity at time zero, and 1 represents the WT intensity at 2 min. In all time courses, n=3 for each time point, data are pooled from three independent experiments; error bars indicate the standard deviation. Statistical significance was determined by unpaired t-test (**C**), by 2-way-ANOVA with Dunnett’s correction (**E** and **H**), or by paired t-test and relates to comparison between ITK-inhibited and non-inhibited cells of the same genotype (**F** and **I**).

To obtain a measure of the highest response observed for each genotype, we determined the normalized fluorescence intensity of cells at the 75^th^ percentile (75-FI). Mutation of Y173 reproducibly reduced the fluorescence intensity at the 75th percentile (Fig 2B), and this difference reached statistical significance at the two-minute time point (Fig 2C). A further, statically significant reduction was observed when Y173F cells were stimulated in the presence of an ITK inhibitor (Fig 2C). Taken together, these results suggest that SLP-76 Y173 facilitates but is not essential for ITK-mediated phosphorylation of PLC-γ1 Y783.

#### An ITK-SLP-76 Y173 axis regulates p-ERK persistence

Further downstream, the PLC-γ1-derived second messenger DAG indirectly triggers ERK phosphorylation [52]. Phospho-ERK is amplified by positive feedback loops that shape a markedly binary response [53–56]. Whereas cells expressing WT SLP-76 exhibited binary accumulation of p-ERK^hi^ cells, SLP-76 deficient J14 cells responded with reduced frequency and markedly reduced intensity of p-ERK staining among the responding cells (Fig 2D).

To capture SLP-76-dependent signaling, we quantified the frequency of strongly responding, p-ERK^hi^ cells (Fig 2E), as defined by the gate shown in Fig 2D. In WT cells, the frequency of p-ERK^hi^ cells peaked at 2-5 minutes and then decayed slowly without fully returning to baseline, over a 30 min time course (Fig 2E). Inhibition of ITK significantly reduced the frequency of p-ERK^hi^ cells at early time points, and reduced the intensity of p-ERK staining among responding cells (Fig 2D-E). Ablation of Y173 similarly reduced the peak frequency of p-ERK^hi^ cells, coupled with a rapid return to a baseline, p-ERK^neg^ state (Fig 2D-E). The Y173F response was further decreased upon inhibition of ITK (Fig 2F), due to reduced intensity of p-ERK staining among responding cells (Fig 2D). Together, these results suggest that SLP-76 Y173 is a prominent downstream target of ITK that promotes the frequency and persistence of the p-ERK^hi^ response, yet additional ITK substrate(s) appear to control the intensity of p-ERK staining among responding cells.

#### p38 phosphorylation strongly depends on ITK and SLP-76 Y173

In addition to ERK, TCR stimulation activates another MAPK family member, p38. Two pathways to TCR-induced activation of p38 have been identified: a conventional, DAG-dependent MAPK cascade, and a supplementary pathway that is driven by LAT-independent, ZAP-70-mediated, direct tyrosine phosphorylation of p38 [57–59]. Published results have not yet assessed the role of SLP-76 in TCR-induced activation of p38. We used phospho-flow to assess the role of SLP-76 in general, and Y173 in particular in mediating the TCR-induced activation of p38.

Following TCR stimulation, we observed distinct phospho-p38 positive and negative populations (Fig 2G). In WT cells, the frequency of p-p38^+^ cells peaked 5-10 min following TCR stimulation and decayed very slowly, with substantial activity still apparent at the 20 min time point (Fig 2G-H). This persistent p-p38 signal contrasted strongly with the rapid decay of p-PLC-γ1 (compare Figs 2H and 2B).

The p38 response was almost completely abrogated in SLP-76-deficient J14 cells (Fig 2G-H), establishing p38 as a strongly SLP-76-dependent signaling event within the TCR pathway. Inhibition of ITK reduced the peak frequency and persistence of p-p38^+^ cells with a rapid return of the cells to the baseline p-p38^neg^ state. A similar reduction in peak frequency and return to baseline was observed in Y173F cells (Fig 2G-H). By the 20 min time point, WT cells had fallen only moderately below their peak response, whereas the majority of ITK-inhibited or Y173F cells had returned close to baseline. Further inhibition of ITK in Y173F cells moderately reduced p-p38, and this difference was statistically significant (Fig 2I). These results suggest that ITK and SLP-76 Y173 are major mediators of the signaling pathways that promote persistent TCR-induced p38 phosphorylation.

Given the involvement of ZAP-70 in TCR-mediated activation of p38, we tested whether Y173 might influence p38 by exerting feedback control over ZAP-70. We indirectly assessed ZAP-70 activity by measuring its direct substrate, phospho-LAT Y171, using a phospho-flow approach. TCR-inducible phosphorylation of LAT was not reduced by the Y173F mutation (Fig 2J-K), suggesting that Y173 exerts signaling effects downstream of ZAP-70.

Taken together, the Y173F mutation reduced the peak amplitude and persistence of three distinct signaling events, phospho-PLCγ1, phospho-ERK, and phospho-p38. All of the above trends were similarly observed when cells were costimulated with TCR/CD28 (Supplementary Fig 1).

### A SLP-76 Y173F mouse model largely phenocopies the conventional developmental phenotypes of ITK^−/−^thymocytes

The cell line studies described above provided evidence that the ITK substrate, SLP-76 Y173, is required to mediate downstream effects of ITK within the TCR signaling pathway. To assess the in vivo significance of the ITK – SLP-76 Y173 signaling axis, we generated a knock in mouse line, in which SLP-76 tyrosine 173 is replaced by phenylalanine (Y173F). To explore the role of Y173 in conventional T cell development, these mice were characterized across multiple parameters, including overall thymic composition, β-selection, and TCR signaling competence. Throughout the study, we noted marked similarities between Y173F and ITK^−/−^thymocytes, which together strengthen the notion that Y173 may serve as an important downstream mediator of ITK.

#### Overall thymic composition of SLP-76 Y173F mice resembles ITK^−/−^

Thymic cellularity was significantly reduced in both young and aged Y173F mice of both sexes (Fig 3A); roughly similar to the reduction that was previously reported in ITK-deficient (ITK^−/−^) mice [29, 30].

**Figure 3.**
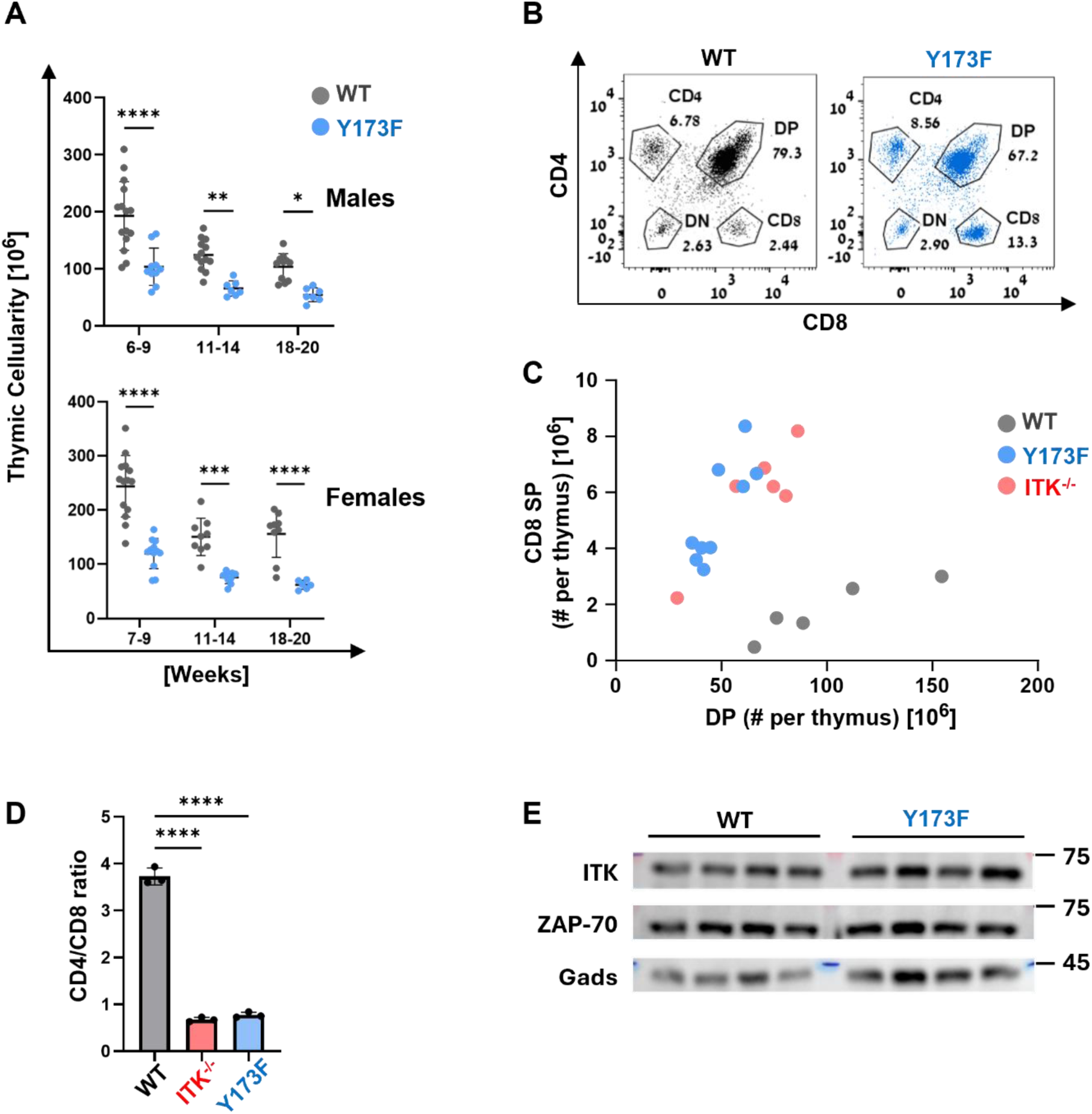
Gross thymic development in SLP-76 Y173F mice resembles ITK^−/−^ thymocytes. (**A**) Total thymocyte cell count in 6-20-week-old male and female mice, either WT (grey) or Y173F (blue). (**B-C**) Thymocytes from 6-9-week-old mice were stained and analyzed by flow cytometry. All results are gated on total live thymocytes. (**B**) Representative thymocyte compartments in WT and Y173F mice. Indicated gates were used to define DN, DP and SP subpopulations. (**C**) 2D plot depicting the absolute numbers of DP and CD8 SP cells. Each dot indicates one 6-9-week-old female mouse of the indicated genotype. (**D**) Ratio of CD4 SP to CD8 SP in a representative experiment. (**E**) Thymocyte lysates from four mice of each genotype were probed by western blot with the indicated antibodies. n≥3 for each genotype. Statistical significance was determined by 2-way-ANOVA with Šídák’s correction (**A**), or one-way-ANOVA with Tukey’s correction (**D**).

Staining for CD4 and CD8 revealed differences in the relative frequencies of each compartment (Fig 3B). Both ITK^−/−^and Y173F mice had fewer DP cells compared to wild type mice, yet a higher frequency and absolute number of CD8 SP cells (Fig 3B and 3C). Consistent with previous reports on ITK^−/−^ mice [30], the increase in CD8 SP thymocytes was associated with a reduced CD4 to CD8 ratio; and this reduction was virtually identical in Y173F and in ITK^−/−^mice (Fig 3D). Given the marked similarity of Y173F and ITK^−/−^ mice, we performed a Western blot to confirm ITK expression in Y173F thymocytes, with Gads and ZAP-70 serving as loading controls. All three proteins were expressed similarly to wild type mice (Fig 3E).

#### β-selection proceeds independently of SLP-76 Y173

The prominent DP compartment suggested that β-selection may proceed independently of SLP-76 Y173. We used two criteria to formally assess β-selection: passage of cells from DN3 to DN4, and allelic exclusion at the TCRβ locus.

To accurately gate on the DN compartment, we used a dump gate to exclude cells bearing CD4, CD8, TCRβ, TCRγδ, and other non-T cell lineage markers, and then used CD44 and CD25 to identify DN1-DN4 (Supplementary Fig 2A). Consistent with intact β-selection, no significant changes were observed in the frequency or absolute number of any of the DN subsets (Supplementary Fig 2B-C).

As an alternative screen for subtle β-selection defects, we considered the possibility that decreased intensity of pre-TCR signaling might impair allelic exclusion at the TCRβ locus, leading to the accumulation of pre-DP cells expressing two distinct TCR Vβ segments. To explore this possibility, we stained thymocytes with V_β_5.1/2 and V_β_8.1/2, while simultaneously staining with a pool of FITC-stained antibodies recognizing other V_β_ gene segments. Since surface TCR expression is very low at the pre-DP stage, we detected V_β_ segments by intracellular staining, while gating on CD3^neg^ CD69^neg^ CD8 cells, which constitute the pre-DP compartment (Supplementary Fig 2D, top panels). Failure of allelic exclusion was defined as the accumulation of cells expressing an additional V_β_ allele along with V_β_5.1/2 or V_β_8.1/2 (Supplementary Fig 2D, bottom panels). This strategy revealed that allelic exclusion was intact in pre-DP Y173F thymocytes (Supplementary Fig 2E). Together, these studies suggest that the pre-TCR signaling pathways that mediate β-selection are largely independent of SLP-76 Y173.

#### Thymic expression of CD5 is profoundly SLP-76 Y173-and ITK-dependent

As an initial assessment of the signaling capacity of Y173F thymocytes, we examined the developmentally regulated expression of CD5. CD5 moderately increases as cells undergo β-selection and further increases upon expression of the full αβ TCR; at which time surface abundance of this marker reflects the intensity of intra-thymic responses to self-peptide-MHC antigens [22, 24].

Despite intact β-selection (Supplementary Fig 2), progression of Y173F thymocytes through DN4 and on to the DP compartment was not accompanied by substantial upregulation of CD5 (Fig 4A-B). Whereas TCR β expression was intact on Y173F DP thymocytes, cell surface CD5 was profoundly reduced (Fig 4C-D), as in ITK^−/−^DP thymocytes (Fig 4E). These results strongly suggest the impairment of TCR responses to self-peptide-MHC in situ.

**Figure 4.**
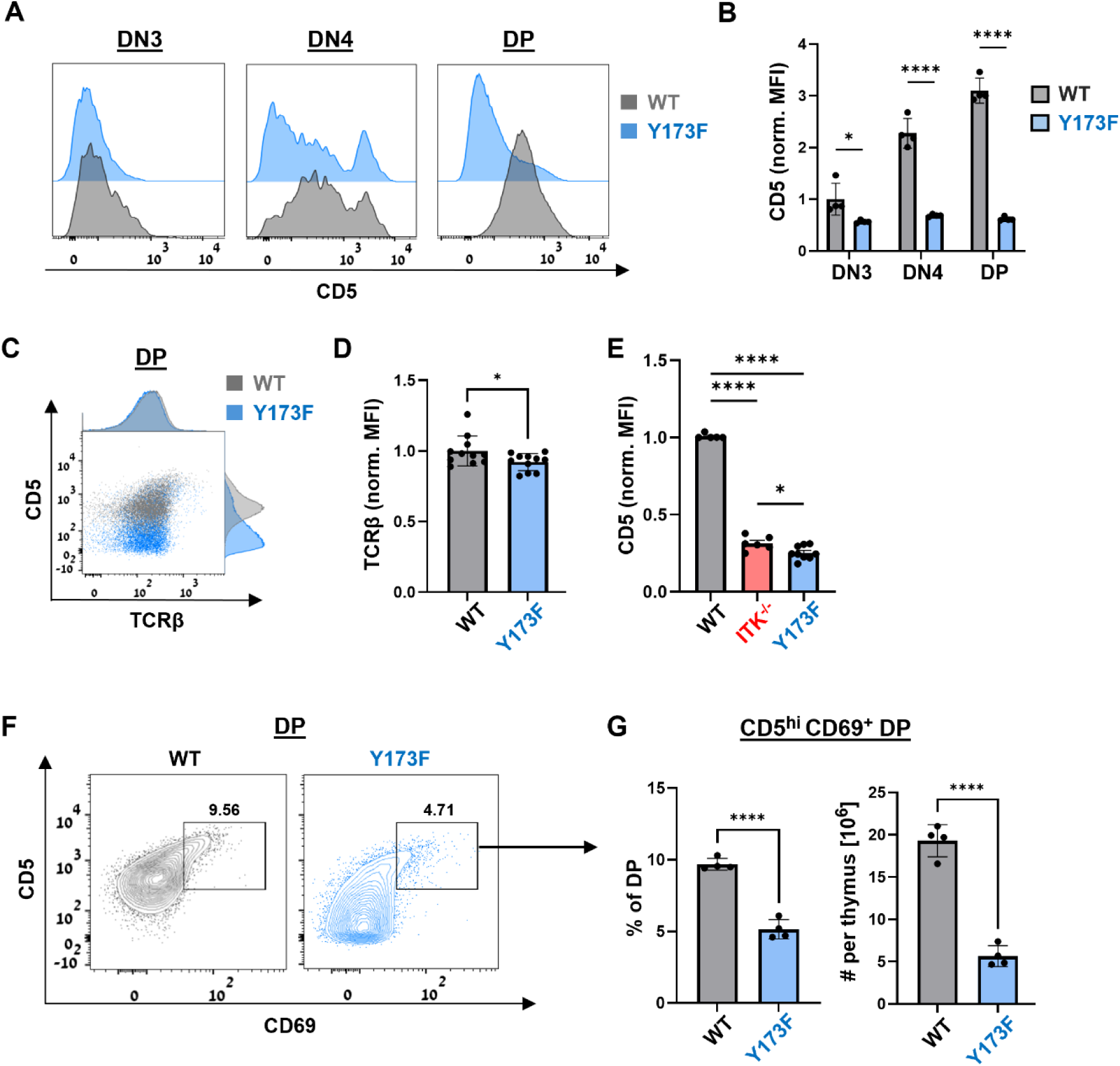
SLP-76 Y173 promotes CD5 expression during and after β-selection. Thymocytes from 6–9-week-old mice were stained and analyzed by flow cytometry, while gating on live DN or DP thymocytes. (**A**) Representative expression of CD5 in DN3 (left), DN4 (middle), and DP (right). When gating on DN populations, we used a dump gate to exclude contaminating macrophages, B cells and DCs, as well as CD4^+^ and CD8^+^ cells. (**B**) Median fluorescence intensity (MFI) of CD5 within each subset, normalized to the average MFI **of** WT DN3. (**C**) Representative expression of TCRβ and CD5 in DP cells. MFI of TCRβ (**D**) and CD5 (**E**) in DP cells, normalized to the average MFI of WT DP. (**F**) Representative expression of CD69 and CD5 in DP. The square gate represents CD69^hi^ DP. (**G**) The frequency (left) and absolute number (right) of CD69^hi^ DP thymocytes. Throughout all plots and bar graphs: WT, grey; Y173F, blue; ITK^−/−^, red. n≥4 for each genotype. Statistical significance was determined by 2-way-ANOVA with Šídák’s correction (**B**), unpaired t-test (**D** and **G**), or one-way ANOVA with Tukey’s correction (**E**).

As cells continue to transit through the DP compartment, positive selection triggers the transient expression of CD69. The overall frequency and number of CD5^hi^ CD69^+^ DP were reduced in Y173F thymocytes (Fig 4F-G), again consistent with a signaling defect. Nevertheless, the presence of a subpopulation of Y173F cells that upregulated CD5 and CD69 (Fig 4F-G) suggests that some cells proceed to positive selection and/or avoid negative selection. This interpretation is consistent with the substantial presence of SP thymic populations (see Fig 3B).

Consistent with positive selection, both CD5 and TCR β increased as Y173F thymocytes progressed to the SP compartments, yet their surface abundance remained significantly lower than in WT SP thymocytes (Supplementary Fig 3). Y173F CD4 SP thymocytes exhibited lower surface expression of TCR b and CD5 (Supplementary Fig 3A-C), closely resembling the phenotype of ITK^−/−^ CD4 SP thymocytes (Supplementary Fig 3D). This tendency was even more apparent in the CD8 SP compartment, and again closely matched the phenotype of ITK^−/−^ CD8 SP thymocytes (Supplementary Fig 3E-H). These data demonstrate that the progression of Y173F thymocytes to the SP compartment is qualitatively different from that exhibited by WT thymocytes, while resembling ITK^−/−^ thymocytes.

#### The ITK–SLP-76 Y173 axis promotes TCR signaling to calcium and MAPK pathways in thymocytes

The profoundly reduced thymic expression of CD5 (Fig 4 and Supplementary Fig 3) is consistent with a requirement for SLP-76 Y173 to mediate thymocyte responsiveness to self-peptide-MHC. To more directly assess TCR-proximal signaling, we stimulated thymocytes ex vivo at 37°C by co-cross-linking CD3, CD4 and CD8, and measured changes in calcium and phospho-proteins, while gating on DP, CD4 SP or CD8 SP thymocytes.

Calcium flux in the DP compartment was generally weak, probably reflecting low TCR expression in this compartment. Nevertheless, a minor population of highly responding cells was observed within the gate shown in Fig 5A, top row. The frequency of highly responding cells during the time frame represented by this gate was substantially reduced by the Y173F mutation (Supplementary Fig 4A).

**Figure 5.**
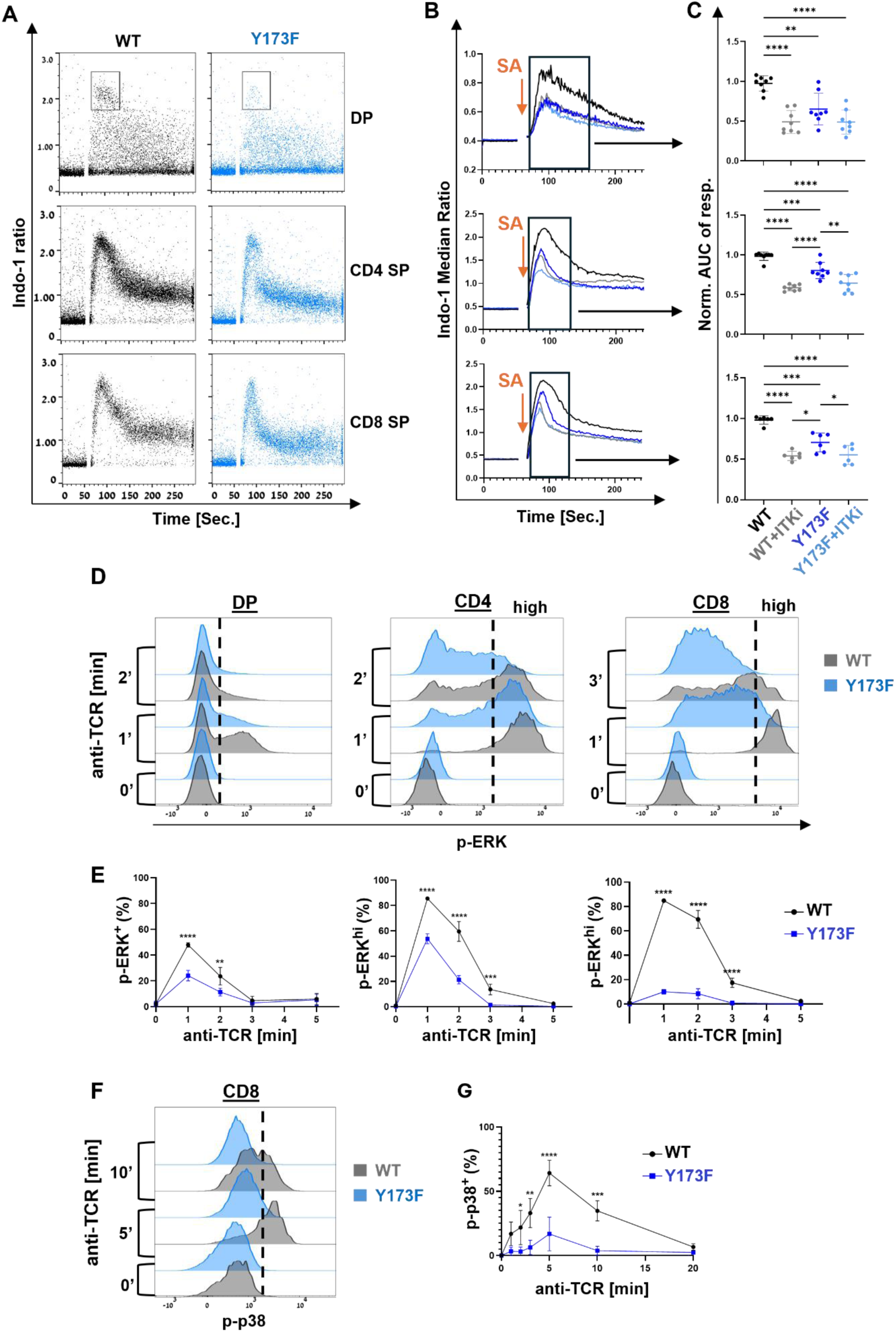
SLP-76 Y173 is required for TCR signaling in thymocytes. Thymocytes from 6-7-week-old mice were differentially barcoded, mixed together, and stained with fluorescent cell surface markers and a mixture of biotinylated stimulatory antibodies (CD3, CD4 and CD8). Cells were preincubated for 15 min at 37°C, in the presence or absence of an ITK inhibitor (BM509744, 3 µM), prior to stimulation. (**A**-**C**) Calcium (Ca^2+^) was measured by flow cytometry at 37°C, using indo-1 as a Ca^2+^ detector. Cells were stimulated by using streptavidin-induced co-crosslinking at the 60 sec time point. (**A**) Representative indo-1 ratio as a function of time, obtained while gating on the indicated compartments. (**B**) Median indo-1 ratio as a function of time while gating on DP (top), CD4 SP (middle), or CD8 SP (bottom). Graphs depict the average of three technical repeats from a single representative experiment. (**C**) Area under the curve (AUC) within the time frame shown in B, normalized to the average area under the curve in WT cells from the same experiment. Data are pooled from three experiments, each including 2-3 technical repeats. (**D-E**) Following stimulation for the indicated time, cells were fixed, permeabilized, and stained intracellularly for phospho-flow. (**D**) Representative p-ERK staining. Within each SP compartment, the high responders gate was set to include 85% of WT cells at the 1 min time point. (**E**) Time course: Frequency of p-ERK^+^ in DP (left), p-ERK^hi^ in CD4 SP (middle), and p-ERK^hi^ in CD8 SP (right). (**F**) Representative p-p38 in CD8 SP. The indicated gate was used to define p-p38^+^ cells. (**G**) Time course: Frequency of p-p38^+^. Data are pooled from three independent experiments; error bars indicate the standard deviation. Statistical analysis by one-way ANOVA with Tukey’s correction (**C**), or 2-way-ANOVA with Šídák’s correction (**E** and **G**).

To probe the involvement of ITK, we repeated the calcium experiment in the presence of an ITK inhibitor. To assess the kinetics of the response, we plotted the median indo-1 ratio as a function of time (Fig 5B, top). The area under this curve (AUC) within the time frame shown in Fig 5B represents the cumulative cellular exposure to high calcium. In DP cells, the AUC was significantly reduced by the Y173F mutation, almost to the extent observed upon inhibition of ITK. Inhibition of ITK in Y173F thymocytes tended to further reduce the AUC, yet the difference between these two data sets did not reach statistical significance (Fig 5C, top). As a control for cell viability and indo-1 loading, no difference in calcium flux was noted between WT and Y173F DP thymocytes when cells were stimulated with Ionomycin (Supplementary Fig 4B).

Within the CD4 SP compartment, calcium flux was initially similar in Y173F and WT cells but decayed more rapidly in Y173F CD4 SP thymocytes (Fig 5A, middle). This reduced persistence was reflected in rapid decay of the median indo-1 ratio (Fig 5B, middle), and markedly reduced AUC (Fig 5C, middle), both of which were further reduced upon inhibition of ITK.

A marked reduction in the persistence of the calcium response was likewise observed in the Y173F CD8 SP compartment (Fig 5A, bottom), resulting in a rapid reduction in the median indo-1 ratio, and markedly reduced AUC, which was further reduced upon inhibition of ITK (Fig B-C, bottom).

Taken together, these results show that SLP-76 Y173 is required to support persistence of the calcium response in DP and SP thymocytes. The similarity of Y173F and ITK-inhibited responses suggests that Y173 is a major component of the pathway by which ITK promotes TCR-induced calcium flux in thymocytes. We observed a trend towards further inhibition of the calcium response upon the addition of an ITK inhibitor in all three compartments, yet this trend reached statistical significance only in the SP populations.

To confirm the presence of a proximal signaling defect in Y173F thymocytes, we performed phospho-flow experiments after co-cross-linking CD3, CD4 and CD8 at 37°C. The weak calcium signal seen in WT DP thymocytes (Fig 5A, top) correlated with weak induction of p-ERK (Fig 5D, left) and an absence of detectable p-p38. Consistent with their calcium signaling defect, Y173F DP thymocytes exhibited markedly reduced induction of p-ERK at the 1 min time point (Fig 5D-E left panels).

Phospho-flow signals were more readily apparent in the SP compartments, where TCR expression is higher. WT CD4 SP thymocytes responded to TCR stimulation with a binary p-ERK signal that peaked at the 1 min time point and returned to baseline by 5 min. Y173F CD4 SP thymocytes exhibited reduced peak frequency and persistence of p-ERK staining (Fig 5D-E, middle panels). An even more marked p-ERK defect was observed in Y173F CD8 SP cells, which exhibited markedly reduced intensity of staining compared to WT (Fig 5D-E, right panels). TCR-induced p38 phosphorylation was most prominent in the WT CD8 SP compartment (Fig 5F). Y173F CD8 SP cells exhibited a delayed response, characterized by profoundly reduced frequency, intensity and persistence of p-p38 staining (Fig 5F-G).

Together, these results suggest that SLP-76 Y173 is required to mediate persistent TCR-induced calcium and MAPK signaling in the thymus, a phenotype that may partially explain the reduced upregulation of CD5.

### An ITK-SLP-76 Y173 axis regulates innate-like thymocyte development

In addition to its role in conventional thymocyte development, ITK profoundly alters the prevalence of innate-like thymic populations. On one hand, ITK promotes the terminal maturation and survival of invariant NKT (iNKT) populations [33–35]. In parallel, ITK limits the development of innate-like SP populations that express PLZF or EOMES [42–44]. To further explore the role of the ITK substrate, SLP-76 Y173, in mediating ITK-dependent developmental phenotypes, we compared innate-like subpopulations in Y173F and ITK^−/−^mice.

#### SLP-76 Y173 is required for the accumulation of thymic DN iNKT1

The term iNKT encompasses several unconventional thymocyte populations, characterized by a semi-invariant αβ TCR that recognizes CD1d-presented lipid antigens. Whereas the earliest iNKT subsets are CD4 SP, some maturation trajectories are accompanied by the loss of CD4 [36]. Differentiation-associated expression of key transcription factors further distinguishes iNKT1, iNKT2 and iNKT17 subsets [60, 61].

Previous studies established that ITK regulates maturation, but not the precursor frequency of iNKT. In particular, ITK^−/−^ mice exhibit a profound decrease in the frequency and number of CD4^neg^ thymic iNKT, accompanied by decreased expression of T-bet within thymic iNKT populations [34]. To explore whether these effects of ITK may depend on ITK-mediated phosphorylation of SLP-76 Y173, we compared iNKT populations in ITK^−/−^ and Y173F mice.

For an initial assessment of the overall frequency and number of thymic iNKT, we gated on TCRβ^+^ CD1dtet^+^ (iNKT) cells in 6–9-week-old mice (Supplemental Fig 5A). This analysis did not reveal statistically significant changes in the frequency or number of total thymic iNKT in ITK^−/−^ thymi. Y173F mice exhibited moderately reduced frequency and number of total thymic iNKT (Supplementary Figs 5A and 5B).

We considered the possibility that changes in the DN iNKT subset may be masked by compensatory changes in CD4 SP iNKT. To bypass this difficulty, we gated based on CD4 expression and added CD44 as a marker of innate-like populations. In WT thymi, CD44^hi^ iNKT populations were clearly detected in both the DN and CD4 SP compartments (Fig 6A, left). The frequency and number of DN iNKT cells were profoundly reduced by the Y173F mutation, yet CD4^+^ iNKT were not affected (Fig 6A-B). Indeed, when compared side by side, both ITK^−/−^ and Y173F mice exhibited marked reductions in the frequency and number of DN iNKT (Fig 6C). This phenotype tended to be more severe in Y173F mice than ITK^−/−^ mice, yet the difference did not reach statistical significance (Fig 6C).

**Figure 6.**
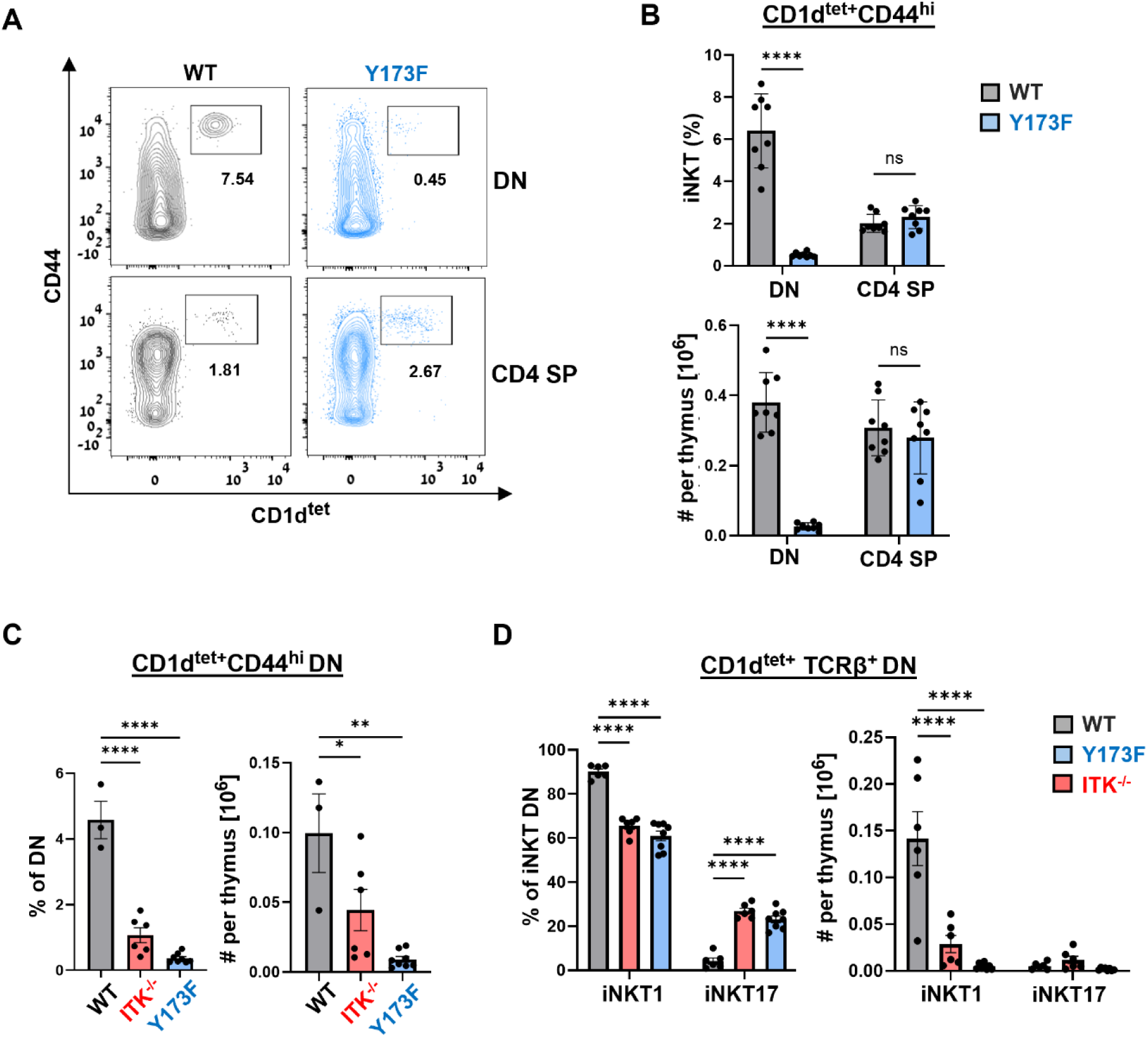
SLP-76 Y173 is required for optimal development or maturation of DN iNKT1. Thymocytes from 6-9-week-old mice were stained and analyzed by flow cytometry, while gating on live DN or CD4 SP cells. (**A**) Representative frequency of CD44^hi^ CD1d-tetramer-positive (iNKT) cells in DN (top) or CD4 SP (bottom). (**B**) Frequency (top) and absolute number (bottom) of iNKT cells within the indicated populations. (**C**) Side by side comparison of the frequency (left) and absolute number (right) of DN iNKT cells in the indicated mouse strains. (**D**) Frequency (left) and absolute number (right) of DN iNKT subsets among DN CD1d^tet+^ TCRβ^+^ thymocytes. The legend relates to all plots and bar graphs: WT, grey; Y173F, blue; ITK^−/−^, red. n≥3 mice for each genotype; error bars indicate the standard deviation. Statistical significance was determined by unpaired t-test (**B**), one-way-ANOVA (**C**), or 2-way-ANOVA (**D**), with Tukey’s correction.

To further explore potential influences of the ITK-SLP-76 Y173F axis on iNKT maturation, we used an established gating strategy [61], to distinguish three iNKT subpopulations: iNKT1, iNKT2, and iNKT17 (Supplementary Fig 5C). All three populations could be distinguished within total thymocytes, but only iNKT1 and iNKT17 were reliably distinguished within the DN population.

ITK^−/−^ and Y173F mice were characterized by decreased frequency and profoundly decreased number of DN iNKT1 cells, accompanied by an increase in the frequency, but not the absolute number of DN iNKT17 cells (Fig 6D). These trends were difficult to distinguish when gating on total iNKT (Supplementary Fig 5D), again suggesting that compensatory changes in the CD4 SP iNKT compartment may mask profound changes in the DN iNKT compartment.

Together, these results confirm and extend the previously described decrease in T-bet expression among ITK^−/−^ iNKT [34], while providing strong evidence that an ITK-SLP-76 Y173 signaling axis is required for the maturation or survival of a CD4^neg^ T-bet^+^ iNKT subset.

#### An ITK-SLP-76 Y173F axis limits the development of PLZF^+^ and EOMES^+^ innate-like SP thymocytes

Multiple innate-like thymic subsets are directly or indirectly limited by ITK, as their prevalence increases in ITK^−/−^ mice. ITK-limited innate-like populations generally express either PLZF or EOMES. A prominent subset includes PLZF^+^ CD4 SP cells that may express either an αβ or γδ TCR. To assess whether the innate-like developmental effects of ITK may depend on its substrate, SLP-76 Y173, we stained for PLZF^+^ and EOMES^+^ innate-like subsets.

Compared to WT, Y173F thymocytes exhibited marked expansion of two PLZF^+^ CD4 SP populations that differed in their expression of TCRβ (Fig 7A). Both increased significantly in frequency and number in Y173F mice (Fig 7B). Consistent with an innate-like phenotype, both populations expressed abundant CD44, a T cell activation marker that characterizes innate-like populations (Fig 7C). In a side-by-side comparison, the increase in frequency was even more dramatic in ITK^−/−^ mice (Fig 7D).

**Figure 7.**
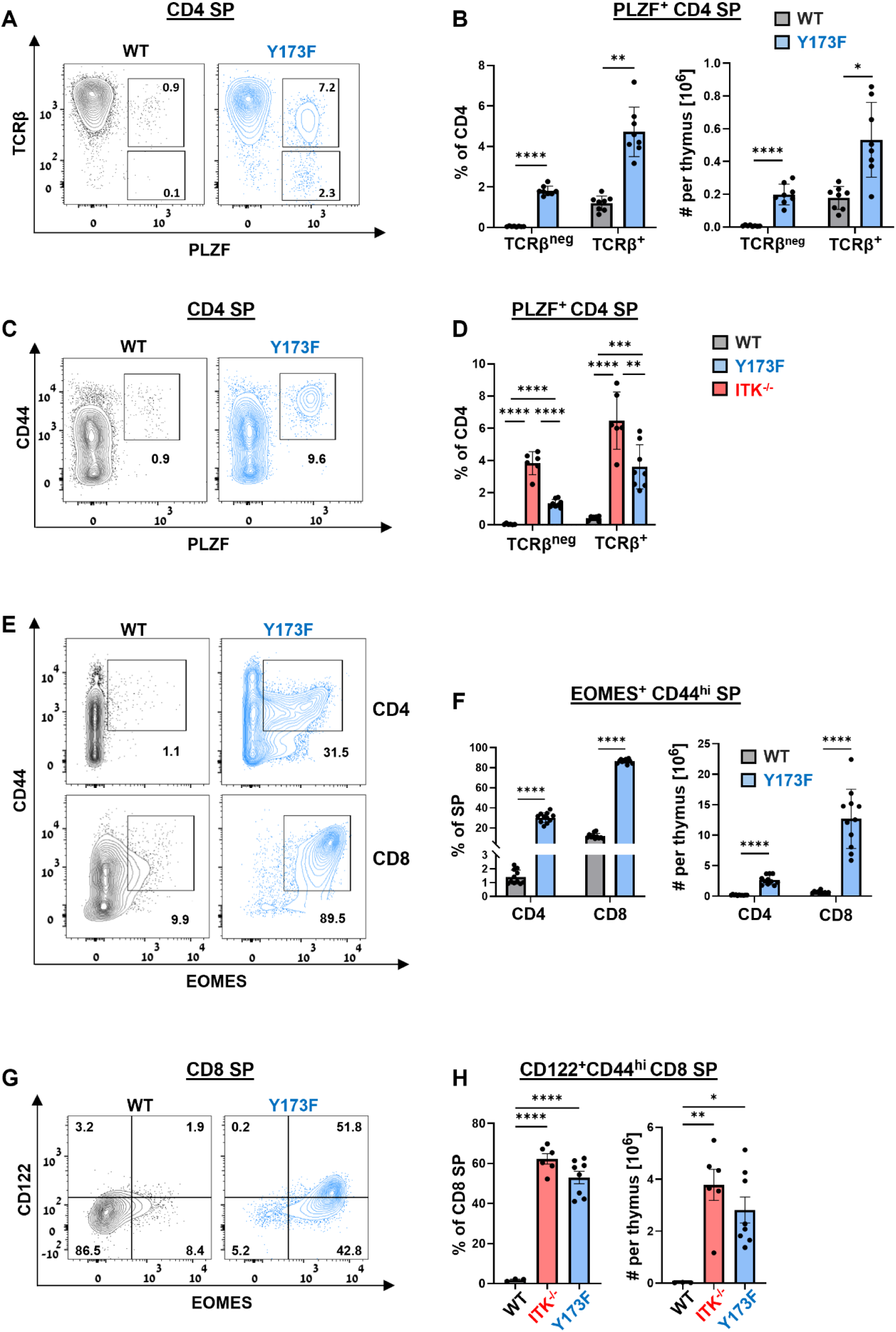
SLP-76 Y173 regulates innate-like SP thymocyte development. Thymocytes from 6-9-week-old mice were stained and analyzed by flow cytometry. (**A**) Representative data depict surface TCRβ and intracellular PLZF, while gating on CD4 SP cells. (**B**) Frequency (left) and absolute number (right) of TCRβ^+^ and TCRβ^neg^ PLZF^+^ populations, as defined in A. (**C**) Representative data depict surface CD44, and intracellular PLZF, while gating on CD4 SP cells. (**D**) Side by side comparison of the frequency of TCRβ^+^ and TCRβ^neg^ PLZF^+^ CD4 SP populations within the indicated mouse strains. (**E-G**) Innate-like populations among TCRβ^+^ SP thymocytes. (**E**) Representative data depict intracellular EOMES, plotted against surface CD44, in CD4 SP (top) or CD8 SP (bottom) cells. (**F**) Frequency (left) and absolute number (right) of the CD44^hi^ EOMES^+^ population in each compartment, as defined in E. (**G**) Representative staining for CD122 and EOMES in CD8 SP cells. (**H**) Side by side comparison of the frequency (left) and absolute number (right) of CD44^hi^ CD122^+^ CD8 SP populations within the indicated mouse strains. Throughout this figure, n≥4 mice for each genotype, data are pooled from at least two independent experiments; error bars indicate the standard deviation. Statistical significance was determined by unpaired t-test (**B** and **F**), or one-way-ANOVA with Tukey’s correction (**D** and **H**).

The prominent TCRβ^neg^ PLZF^+^ CD4 SP population seen in Fig 7A most likely encompasses γδ^+^ CD4 SP thymocytes. To test this interpretation, we examined the frequency of γδ T cells in Y173F thymi. The Y173F mutation did not affect the frequency or the number of DN γδ^+^ thymocytes (Supplementary Fig 6 A-B) but markedly increased the frequency and number of γδ T cells observed after excluding the DN compartment (Supplementary Fig 6 C-D). Since non-DN γδ thymocytes generally express CD4, we interpret these results to indicate an increase in γδ^+^ PLZF^+^ CD4 SP population within the Y173F thymus.

A distinct, unusual population of CD44^hi^ EOMES^+^ CD4 SP thymocytes is found in ITK^−/−^ thymocytes [41], which also exhibit a prominent population of CD44^hi^ EOMES^+^ CD8 SP. Both populations markedly increased both in frequency and in number in Y173F mice (Fig 7E-F). Consistent with an innate-like phenotype, Y173F EOMES^+^ CD8 SP showed an increased tendency to co-express CD122 (Fig 7G), For side-by side comparison of innate-like CD8 populations in Y173F and ITK^−/−^ mice, we compared the prevalence of CD44^hi^ CD122^+^ CD8 SP thymocytes (Supplementary Fig 6E). Both the frequency and number of this population increased dramatically in Y173F and ITK^−/−^ mice (Fig 7H).

Taken together, our results demonstrate remarkable similarity between ITK^−/−^ and Y173F mice over multiple innate like populations, providing strong evidence that ITK-mediated phosphorylation of SLP-76 Y173 is required to mediate the phenotypic effects of ITK throughout thymic development.

## Discussion

As the third member of the TCR-induced tyrosine kinase cascade, ITK occupies a well-defined niche within the TCR signaling pathway. By phosphorylating PLC-γ1, ITK directly promotes the production of second messengers that are required for TCR responsiveness. Cell line studies provide strong evidence that Impaired PLC-γ1 phosphorylation is likely to mediate many of the developmental phenotypes of ITK^−/−^ mice [6, 7]. Yet this compelling hypothesis has never been tested in a mouse model, as PLC-γ1^−/−^ mice have an embryonic lethal phenotype and PLC-γ1 Y783F mice have not been reported.

To directly test the implications of limiting thymic PLC-γ1 activation, two groups independently characterized LAT Y136F mice, in which a point mutation specifically impairs the recruitment of PLC-γ1 to the LAT signalosome [62, 63]. ITK^−/−^ and LAT Y136F mice share impaired PLC-γ1 phosphorylation [5, 63]; yet their thymic phenotypes are distinct. Whereas ITK^−/−^ mice exhibit a subtle impairment of β-selection accompanied by moderately decreased thymic cellularity and a decreased ratio of CD4 to CD8 SP [29–31]; LAT Y136F mice exhibit profoundly impaired β-selection, a 10-fold reduction in thymic size, and a markedly increased ratio of CD4 to CD8 SP [62, 63]. Downstream of PLC-γ1, DAG binds and activates RAS-GRP, triggering MAPK signaling [52].

Despite moderately impaired β-selection and intact expression of CD5 on RasGRP1^−/−^ DP thymocytes [64, 65], positive selection is markedly impaired, resulting in reduced generation of CD4 and CD8 SP thymocytes [66]. A profound reduction in total iNKT cells in RasGRP1^−/−^ thymocytes is associated with a selective loss of CD4^+^ iNKT [67]. These phenotypes are distinct from ITK^−/−^ mice, which exhibit low expression of CD5 on DP thymocytes [30, 31], coupled with moderately impaired positive selection [29, 30, 32] and reduced numbers of CD4^neg^ iNKT thymocytes [33, 34].

The relatively mild phenotypes of ITK^−/−^ mice may reflect partial impairment of PLC-γ1 phosphorylation, due to compensatory activity of RLK, yet even RLK^−/−^ITK^−/−^ double-deficient mice exhibit thymic phenotypes that are distinct and less severe than either LAT Y136F or Ras-GRP^−/−^ mice [30]. Based on these considerations, we propose that PLC-γ1 phosphorylation may be necessary, but not sufficient to fully mediate the signaling and developmental functions of ITK. Just as ZAP-70 phosphorylates at least seven known substrate sites - four on LAT and three on SLP-76 – all of which are required for TCR signal transduction [9, 10, 68–70]; we propose that additional substrates of ITK may be required to mediate its downstream effects.

In this study, we provide evidence that SLP-76 Y173 is required to mediate the signaling and developmental effects of ITK. At short time scales, Y173 was required for ITK-dependent signaling events, both in T cell lines and in primary thymocytes. At longer time scales, the thymocyte developmental requirements for Y173 closely mimicked the developmental effects of ITK. Together, these lines of evidence establish SLP-76 Y173 as an obligatory mediator of the biological roles of ITK.

One goal of this study was to better define the short-term signaling functions of SLP-76 Y173. Previous functional analyses highlighted the role of Y173 in mediating TCR- and FcεRI-induced phosphorylation of PLCγ1 [16]. The mechanistic basis for Y173-dependent PLC-γ1 phosphorylation remained unclear, as Y173 is not required for ITK activity per se, nor was it required for the TCR-inducible association of SLP-76 with ITK, Vav, Nck or PLC-γ1, as measured by co-immunoprecipitation [16, 17]. Given intact SLP-76 complex formation and intact ITK activation, how might the Y173F mutation impair downstream signaling?

A cell-free reconstitution experiment suggested a plausible mechanism, as inclusion of a SLP-76 p-Y173 peptide increased ITK-mediated phosphorylation of PLC-γ1 in vitro [71]. This result suggested that docking of the C-terminal SH2 domain of PLC-γ1 onto phospho-Y173 may render it more accessible to ITK kinase activity. An important caveat to this interpretation is that Y173 was not required for TCR-induced accumulation of phospho-PLC-γ1 within the SLP-76-nucleated complex [17]. Since active PLC-γ1 may be released from LAT [72], the above observations may instead support a model in which p-Y173 directly or indirectly promotes the flux of PLC-γ1 through the LAT complex leading to the accumulation of phospho-PLC-γ1 outside of the LAT signalosome.

To test this interpretation, we adopted a phospho-flow approach to assess total phospho-PLC-γ1 on a single cell basis. We found that the Y173F mutation moderately reduced peak p-PLC-γ1 without reducing p-LAT Y171. These observations support the notion that Y173 acts downstream of LAT to promote the accumulation of p-PLC-γ1. Yet PLC-γ1 phosphorylation had a substantial Y173-independent component, as evidenced by the further reduction in p-PLCγ1, when Y173F cells were stimulated in the presence of an ITK inhibitor.

Active PLC-γ1 produces IP3, which triggers a rapid increase in intracellular calcium. As expected, both the amplitude and the persistence of the TCR-induced calcium were markedly reduced when SP thymocytes were stimulated in the presence of an ITK inhibitor. The Y173F mutation markedly blunted both the amplitude and the persistence of calcium flux, but these parameters were further reduced when Y173F SP thymocytes were stimulated in the presence of an ITK inhibitor. When considered together with the p-PLC-γ1 phospho-flow data, these results provide strong evidence that SLP-76 Y173 facilitates but is not absolutely required for TCR-induced accumulation of phospho-PLCγ1, both in cell lines and in primary thymocytes.

Our data are consistent with the docking model suggested by Devkota [71], but do not exclude alternative mechanisms. The substantial ITK-dependent, but Y173-indpendent components of PLC-γ1 phosphorylation and calcium flux suggest that docking of PLC-γ1 on p-Y173 is not absolutely required to bridge its interaction with ITK. In addition to the docking model, we consider the possibility that Y173 may facilitate the accumulation of cellular p-PLC-γ1, by promoting the release of active PLC-γ1 from the LAT signalosome, or by increasing the lifetime of p-PLC-γ1 molecules.

Downstream of PLC-γ1, TCR signaling synergizes with CD28 to induce NF-κB activity. In contrast to its partial effect on PLC-γ1, the Y173F mutation profoundly impaired TCR/CD28-induced nuclear translocation of c-Rel, an NF-κB family member [17]. In side-by-side experiments, the Y173F mutation strongly inhibited NF-κB-dependent luciferase reporters while not significantly reducing the activity of an NFAT-luciferase reporter [17]. In a similar fashion, inhibition of ITK disproportionately impaired the nuclear translocation of the NF-κB family member, p65/RelA, under conditions where NFAT was not substantially affected [73]. Together, these results suggest heightened sensitivity of the NF-κB pathway to upstream ITK activity; alternatively, the results may reflect a proposed direct effect of SLP-76 on the nuclear pore transport of NF-κB family members [74].

To distinguish between these possibilities, we asked whether SLP-76 Y173 affects signaling events just upstream of NF-κB nuclear translocation. Consistent with an upstream signaling defect, we found that Y173 was required to trigger the phosphorylation and degradation of IκB, and for the consequent nuclear translocation of RelA. Further downstream, Y173 was required to induce the transcription of c-Rel and IL-2. This result is consistent with an influence on NFAT or NF-κB, factors that are required for the inducible transcription of c-Rel [75, 76] and IL-2 [77].

Remarkably, Nur77 transcription was SLP-76-dependent but Y173-independent. This result validates the structural and functional integrity of the Y173F protein, while supporting a pathway-specific requirement for this site. Nur77 is an orphan family transcription factor whose expression tightly correlates with the strength of TCR signaling. Nur77 transcription is largely NFAT-independent, and only modestly affected by the MAPK pathway [78]. Similar to Y173, ITK catalytic activity is selectively required for antigen-induced transcription of IL-2 [73], but not of Nur77 [73, 79]. This intriguing correlation first suggested to us that an ITK-Y173 axis may control particular aspects of T cell activation downstream of the TCR.

Downstream of the TCR, positive and negative feedback loops control the amplitude and persistence of TCR-proximal signaling events [53, 54, 80, 81]. The contrast between a partial requirement for Y173 to mediate PLC-γ1 phosphorylation at two min and the strong requirement for NF-κB activation at twenty min, suggests a potential role for Y173 in sustaining and propagating early TCR signaling events to later time points. Supporting this notion, both Y173 and ITK were required to support the peak amplitude of p-ERK and p-p38, and for the persistence of this signal as late as thirty min after TCR stimulation. Y173 was likewise required to support ERK and p38 phosphorylation in TCR-stimulated SP thymocytes.

To the best of our knowledge, this study is the first to conclusively show a role of SLP-76 and ITK in TCR-induced p38 activation. Previous studies showed that SLP-76 in general and Y173 in particular are required to mediate p38 phosphorylation in other ITAM-initiated signaling pathways, including FcεRI-induced phosphorylation of p38 in mast cells [16]; moreover, SLP-76 is required for p38 activation in integrin-activated neutrophils [82]. Consistent with our results, a previous study found reduced persistence of TCR-induced phospho-p38 in expanded ITK^−/−^ CD8 T cell blasts, by semi-quantitative, western blot data [83]. Our quantitative phospho-flow data strongly support the requirement for an ITK-SLP-76 Y173 axis to mediate sustained p38 phosphorylation in the TCR signaling pathway.

Together, these results suggest that Y173 facilitates the sustainment of TCR-initiated events over time, both in human T cells and in mouse thymocytes. By extension, these findings suggest that NF-κB activation may depend on sustained TCR signaling. Further research will be required to identify the mechanisms by which SLP-76 phospho-Y173 promotes TCR signal persistence.

To explore the role of Y173 in mediating long-term biological effects of ITK, we compared the thymic phenotypes of Y173F and ITK^−/−^ mice. Y173F mice resembled ITK^−/−^ mice over multiple conventional phenotypes, including reduced thymic cellularity, altered subset composition, and reduced upregulation of the CD5 activation marker. Further remarkable similarities were observed in the development and differentiation of unconventional thymocyte populations. ITK and SLP-76 Y173 were similarly required to promote the differentiation and accumulation of a DN iNKT1 population, and to limit the accumulation of innate-like PLZF^+^ CD4 SP cells and EOMES^+^ CD4 SP and CD8 SP populations. Among these phenotypes, the expansion of an EOMES^+^ CD4 SP population is associated with alterations in the thymic cytokine environment, as observed following the loss of ITK protein or activity [41, 61]. The expansion of this population in Y173F thymocytes provides particularly strong evidence that ITK-mediated phosphorylation of Y173 is required for optimal TCR signaling and for limiting the expansion of innate-like populations.

The Y173F phenotypes were likewise remarkably similar to the phenotypes of SLP-76 Y145F mice [40, 41, 45]. Despite their location on the same signaling protein, SLP-76 Y145 and Y173 have mechanistically distinct roles in the TCR signaling pathway. Y145 is phosphorylated by ZAP-70 and is one of the sites that are required to recruit and activate ITK. In contrast, we have shown that SLP-76 Y173 is not required for the activation of ITK but is one of its downstream substrates. Taken together, the broad phenotypic similarities of SLP-76 Y145F, ITK^−/−^and SLP-76 Y173F mice suggest a coordinated signaling axis, in which Y145-mediated recruitment of ITK leads to ITK-mediated phosphorylation of Y173, which activates key molecular mechanisms that control developmental phenotypes. These mechanisms may be mediated, in part by phosphorylation of PLC-γ1 and may also involve the recruitment of a yet unidentified signaling protein to p-Y173. Either way, the phenotypes described here establish SLP-76 Y173 as an obligatory mediator of the in vivo biological roles of ITK.

## Acknowledgments

This research was supported in part by the Intramural Research Program of the National Institutes of Health (NIH), and in part by a grant from the United States - Israel Binational Science Foundation (BSF grant # 2021035). The contributions of the NIH authors are considered Works of the United States Government. However, the findings and conclusions presented in this paper are those of the authors and do not necessarily reflect the views of the NIH or the U.S. Department of Health and Human Services. Additional support was provided a grant to DY from the Israel Science Foundation (Grant #283/22), by the Minerva Stiftung Center Program on Cell Intelligence and by the Colleck Research Fund. We thank the NIH Tetramer Core Facility (NIH Contract 75N93020D00005 and RRID:SCR_026557) for providing PBS-57 loaded and unloaded CD1d-tetramers. Ofer Sheker, Amir Grau, Rotem Honen Kadosh and Jeeda Bisharat from the Biomedical Core Facility (BCF) of the Rappaport Faculty of Medicine, provided access to flow cytometry equipment and excellent technical support. We thank Prof. Yotam Bar-On for providing the TCR V_β_ screening kit. We thank Julie Reilley and Selamawit Belda from the NIH for excellent technical assistance. We thank the chief veterinarian of the Technion Preclinical Authority, Dr. Dvir Mintz, as well as Paul Zannou, Igor Kosarev, and the rest of the excellent veterinary staffs of the Technion and the NIH for professional assistance with the care, treatment and housing of the mice.

