## Supplemental Figures for "An ITK – SLP-76 Y173 Axis Regulates TCR Signal Persistence and Thymic Development"

#### Supplementary Data

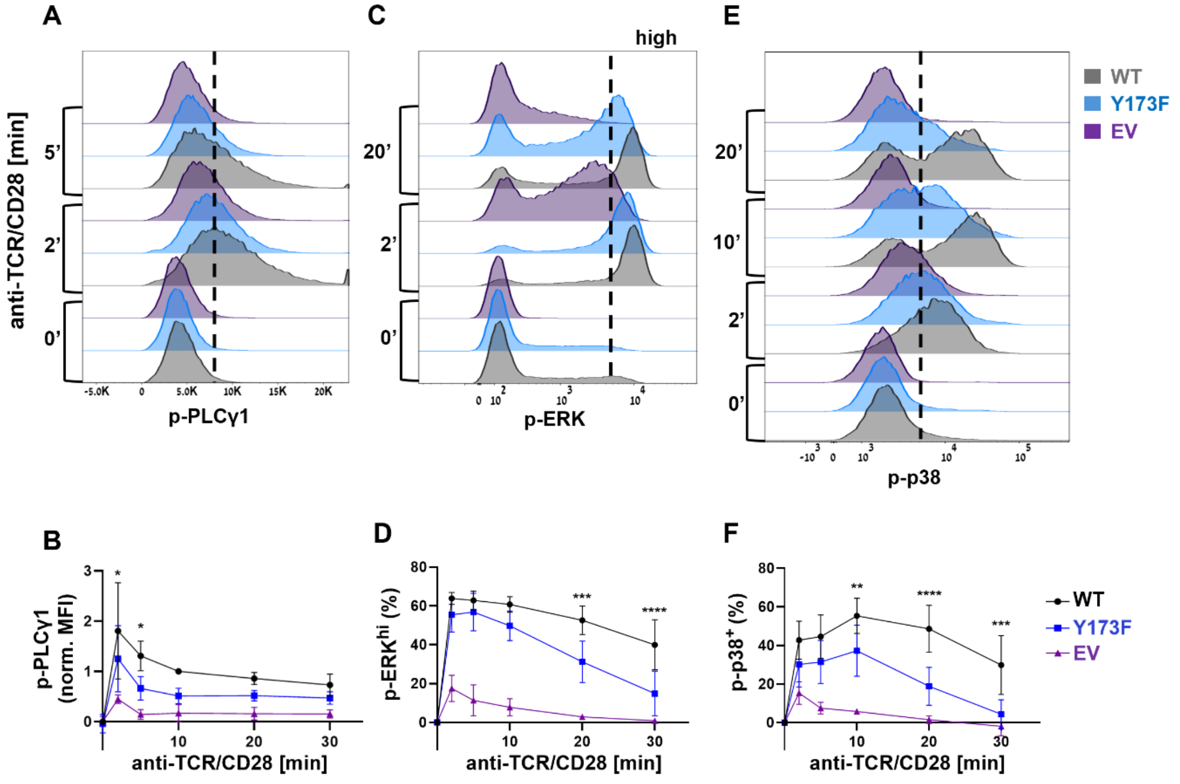


**Supplementary Figure 1. SLP-76 Y173 promotes the persistence of TCR/CD28 signaling.** The indicated cell lines were differentially barcoded and mixed together, then incubated for 15 min at 37°C, prior to stimulation with soluble anti-TCR (C305, 1:1000) and anti-CD28 (clone 28.2, 2 µg/ml). At the indicated time points, cells were fixed, permeabilized, and stained intracellularly for phospho-flow. (**A**) Representative p-PLC-γ1. (**B**) Time course: Min/max normalized median fluorescence intensity (MFI) of p-PLC-γ1. Zero represents the WT intensity at time zero, and 1 represents the average WT intensity at 10 min. (**C**) Representative p-ERK. The high responders gate was set to include 60% of WT cells at the 10 min time point. (**D**) Time course: Frequency of p-ERK$^{hi}$. (**E**) Representative p-p38. The indicated gate was used to define p-p38$^{+}$ cells. (**F**) Time course: Baseline-subtracted frequency of p-p38$^{+}$. $n \geq 3$ for each time point, data are pooled from three independent experiments; error bars indicate the standard deviation. Statistical analysis by 2-way-ANOVA with Dunnett’s correction.

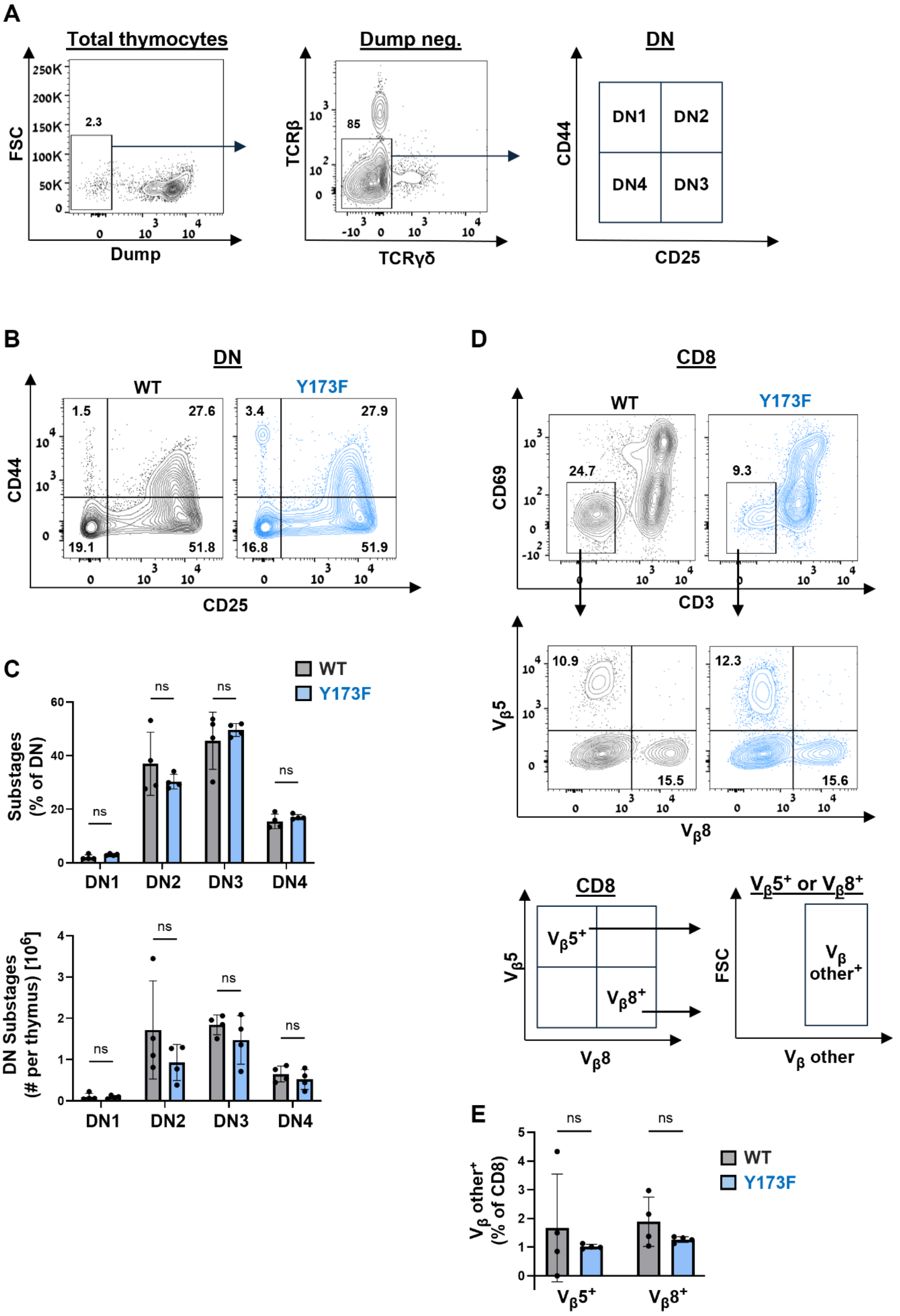
A
Total thymocytes
FSC
Dump
2.3
Dump neg.
TCRβ
TCRγδ
85
DN
CD44
CD25
DN1
DN2
DN4
DN3
B
DN
WT
Y173F
1.5
27.6
19.1
51.8
3.4
27.9
16.8
51.9
CD44
CD25
C
WT
Y173F
Substages (% of DN)
DN Substages (# per thymus) [10^6]
ns
DN1
DN2
DN3
DN4
D
CD8
WT
Y173F
CD69
CD3
24.7
9.3
Vβ5
Vβ8
10.9
15.5
12.3
15.6
CD8
Vβ5+
Vβ8+
Vβ5+ or Vβ8+
FSC
Vβ other+
Vβ other
E
Vβ other+ (% of CD8)
WT
Y173F

**Supplementary Figure 2. SLP-76 Y173 did not detectably influence β-selection or allelic exclusion.** Thymocytes from 7-9-week-old mice were stained and analyzed by flow cytometry. (**A**) Gating strategy. To focus on β-selection, a dump gate was first used to exclude $CD8^+$ and $CD4^+$ cells as well as contaminating macrophages, B cells and dendritic cells, then specific stains were used to exclude $TCR\beta^+$ and $\gamma\delta^+$ cells. (**B**) Representative staining for DN sub-compartments (DN1-4), within the $TCR\beta^{neg}$ $\gamma\delta^{neg}$ DN population, as defined in A. (**C**) Frequency (top) and absolute number (bottom) of each DN sub-compartment. (**D**) Gating strategy for allelic exclusion. Pre-DP cells were defined as $CD3^{neg}$ $CD69^{neg}$ CD8 SP (top). Identification of $V_\beta5^+$ or $V_\beta8^+$ pre-DP (middle). Cells co-staining for any of a mixture of TCR Vβ alleles ($V_\beta$ other) were identified within the $V_\beta5^+$ and $V_\beta8^+$ pre-DP populations (bottom). (**E**) Frequency of $V_\beta5^+$ $V_\beta other^+$ and $V_\beta8^+$ $V_\beta other^+$ pre-DP cells, as defined in D. The legend relates to all plots and graphs: WT, grey; Y173F, blue. n=4 mice for each genotype; error bars indicate the standard deviation. Statistical significance was determined by 2-way-ANOVA with Šídák's correction (**C**), or unpaired t-test (**E**).

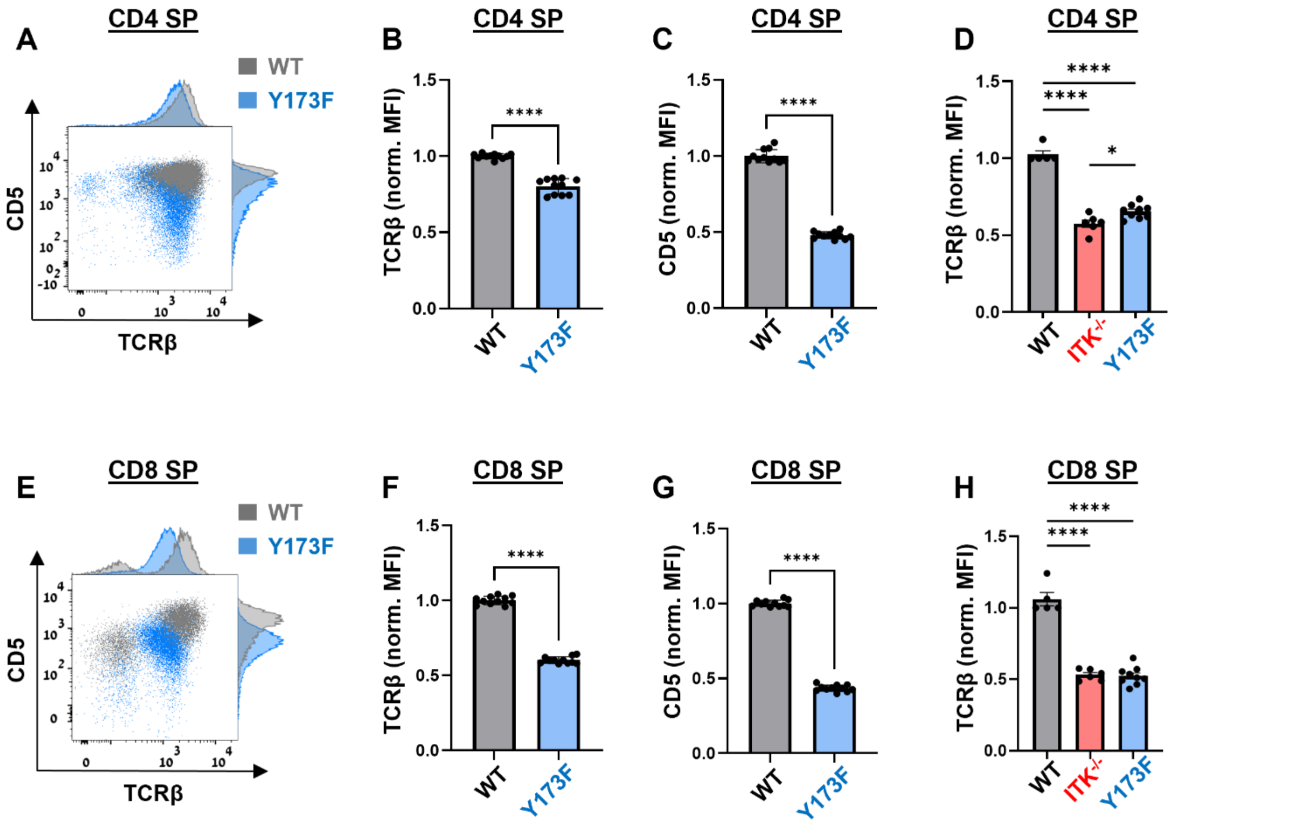


**Supplementary Figure 3. The Y173F mutation impairs CD5 expression in the SP compartments.** Thymocytes from 6-9-week-old mice were stained and analyzed by flow cytometry, while gating on live CD4 (**A**-**D**) or CD8 (**E**-**H**) SP cells. (**A** and **E**) Representative expression of TCRβ and CD5 in the indicated SP compartments. (**B** and **F**) Median fluorescence (MFI) of TCRβ, normalized to average TCRβ expression in WT. (**C** and **G**) Median fluorescence (MFI) of CD5, normalized to average CD5 expression in WT. (**D** and **H**) Side by side comparison of the normalized MFI of TCRβ in the indicated mouse strains and compartments. The legend relates to all bar graphs and dot plots: WT, grey; Y173F, blue; ITK$^{-/-}$, red. n≥5 for each genotype, data are pooled from at least three independent experiments; error bars indicate the standard deviation. Statistical significance was determined by unpaired t-test (**B**-**C**, **F**-**G**), or one-way ANOVA with Tukey's correction (**D** and **H**).

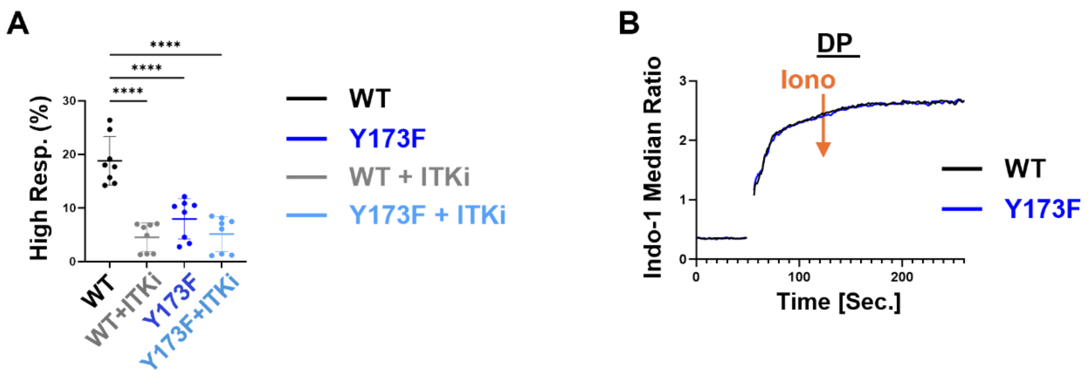


**Supplementary Figure 4. Supplementary calcium data.** (**A**) Calcium flux was measured by FACS at 37°C in indo-1-loaded, differentially barcoded, streptavidin-stimulated thymocytes, exactly as in Fig 5A, in the presence or absence of an ITK inhibitor (BM509744, 3 µM). Shown is the frequency of high-responding DP cells within the gate shown in Fig 5A. (**B**) Instead of streptavidin, cells were stimulated at the 60 sec time point with 1 µM ionomycin. The median indo-1 ratio is presented as function of time while gating on DP. This panel depicts the average of three technical repeats from a single representative experiment. Data are pooled from three experiments, each including two to three technical repeats. Error bars indicate the standard deviation. Statistical analysis was by 2-way-ANOVA with Šídák's correction.

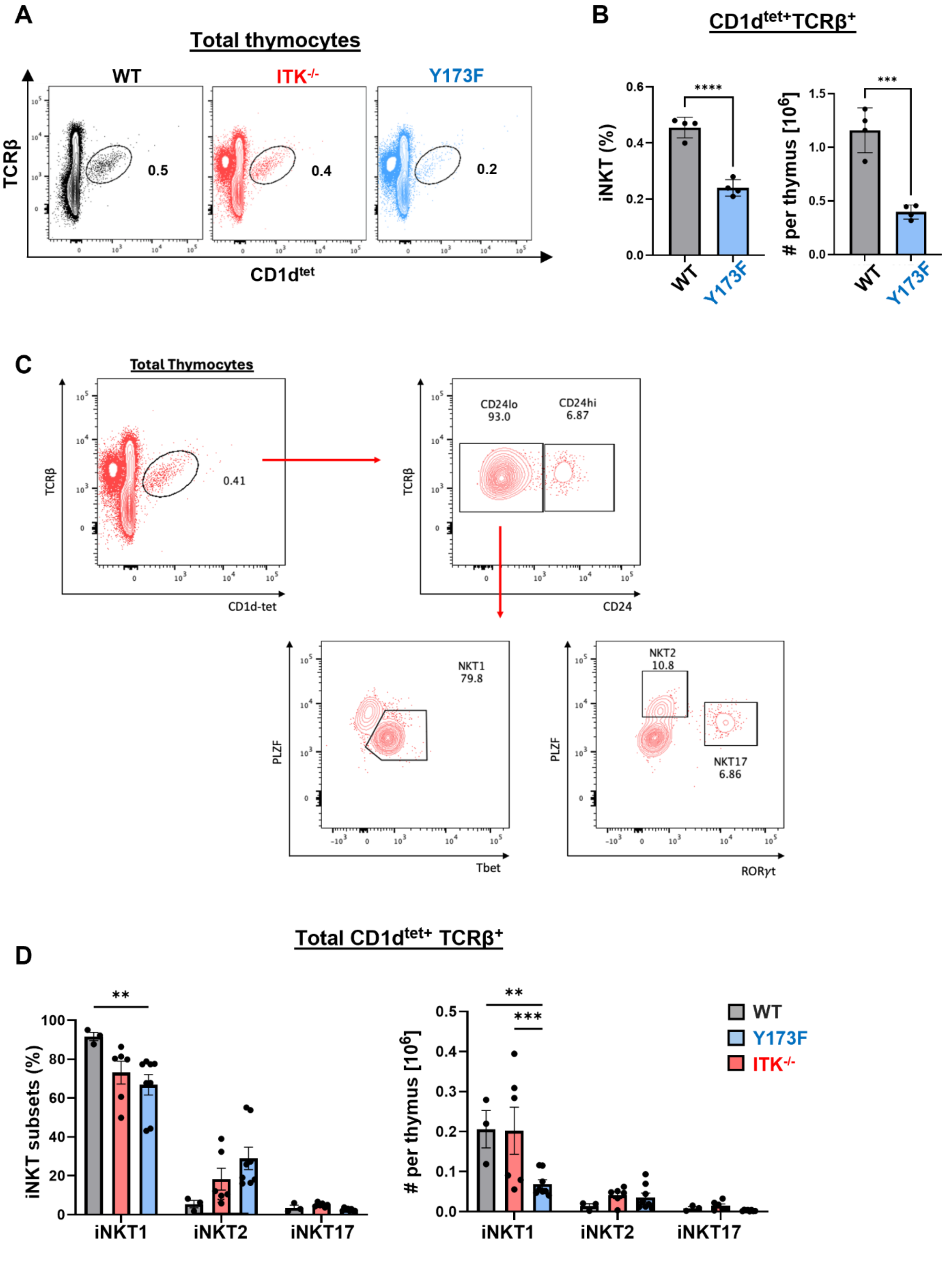
A
Total thymocytes
WT
ITK-/-
Y173F
TCRβ
CD1dtet
0.5
0.4
0.2
B
CD1dtet+TCRβ+
iNKT (%)
### per thymus [10^6]
****
***
WT
Y173F
C
Total Thymocytes
TCRβ
CD1d-tet
0.41
CD24lo
93.0
CD24hi
6.87
CD24
NKT1
79.8
PLZF
Tbet
NKT2
10.8
NKT17
6.86
RORγt
D
Total CD1dtet+ TCRβ+
iNKT subsets (%)
### per thymus [10^6]
iNKT1
iNKT2
iNKT17
**
***
WT
Y173F
ITK-/-

**Supplementary Figure 5. Total thymocyte iNKT data.** Thymocytes from 6-9-week-old mice were stained and analyzed by flow cytometry, while gating on total live thymocytes. (**A**) Representative frequency of CD1d-tetramer$^{+}$ TCRβ$^{+}$ (iNKT) cells. (**B**) Frequency (left) and absolute number (right) of iNKT cells within the indicated mouse strains. (**C**) iNKT subset gating strategy, depicting representative data from an ITK$^{-/-}$ mouse. (**D**) Frequency (left) and absolute number (right) of iNKT subsets within the indicated mouse strains. The legend relates to all plots and bar graphs: WT, grey; Y173F, blue; ITK$^{-/-}$, red. n≥3 mice for each genotype; error bars indicate the standard deviation. Statistical significance was determined by one-way-ANOVA with Tukey's correction (**B**), or 2-way-ANOVA with Šídák's correction (**D**).

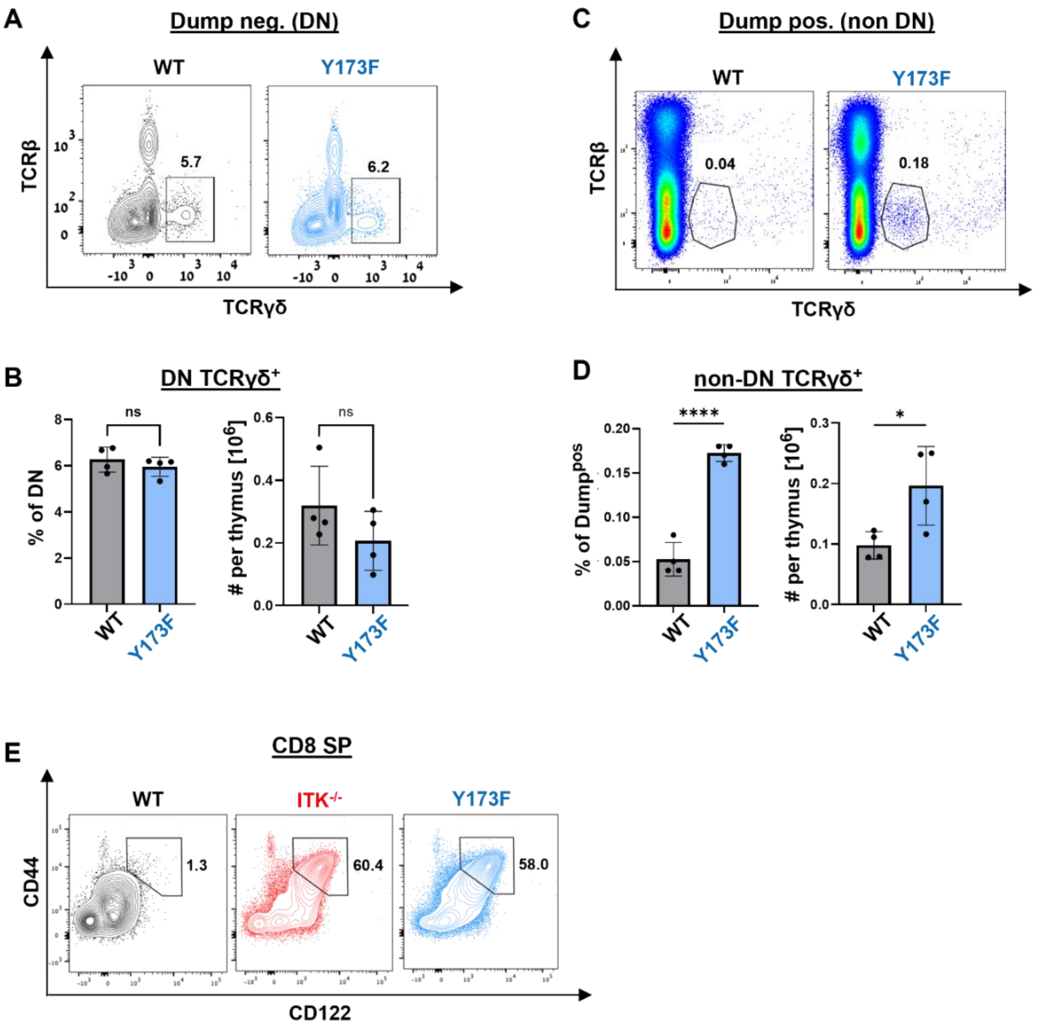


**Supplementary Figure 6. Supplementary innate-like development data.** Thymocytes from 7-9 week-old mice were stained and analyzed by flow cytometry, while gating on live cells. (**A**-**D**) A dump gate was used to exclude $CD8^+$ and $CD4^+$ cells as well as contaminating macrophages, B cells and dendritic cells, as in Supp. Fig 2A, left. (**A**) Representative results. The indicated gate was used to define $TCR\gamma\delta^+$ DN cells. (**B**) Frequency (left) and absolute number (right) of $TCR\gamma\delta^+$ DN cells. (**C**) Representative $dump^+$ cells, a designation that includes all but the DN population. The indicated gate was used to define $TCR\gamma\delta^+$ non-DN cells. (**D**) Frequency (left) and absolute number (right) of $TCR\gamma\delta^+$ non-DN cells. (**E**) Representative staining CD44 and CD122 while gating on CD8 SP. The indicated gate was used to define innate-like CD8 SP in Fig 7H. The legend relates to all plots and graphs: WT, grey; Y173F, blue, $ITK^{-/-}$, red. n=4 mice for each genotype; error bars indicate the standard deviation. Statistical significance was determined by unpaired t-test.
